# High-fat diet ablates an insulin-responsive pool of GLUT4 glucose transporters in skeletal muscle

**DOI:** 10.1101/2025.06.29.662135

**Authors:** Youjia Hu, Stacey N. Brown, Ali Nasiri, Don T. Li, Haiyan Wang, Gregory D. Cartee, Gerald I. Shulman, Jonathan S. Bogan

**Author notes:** Department of Pediatric Orthopedics, Hospital for Special Surgery, New York, NY 10021, USA.

## Abstract

To stimulate glucose uptake in muscle, insulin mobilizes GLUT4 glucose transporters to the cell surface. During fasting, GLUT4 and the transmembrane aminopeptidase IRAP are trapped in intracellular, insulin-responsive vesicles bound by TUG, AS160, and Usp25m proteins. Here we show that Usp25m, a protease, is required for the bulk of insulin-stimulated TUG cleavage and consequent vesicle mobilization and glucose uptake. Efficient TUG cleavage also requires AS160. In mice with diet-induced insulin resistance, Usp25m abundance is reduced, IRAP is mislocalized during fasting, and TUG cleavage is impaired; effects of Usp25m and TUG deletion to alter insulin-stimulated and fasting glucose uptake, respectively, are ablated. We conclude that skeletal muscle insulin resistance results in part from altered membrane trafficking of GLUT4 and IRAP during fasting. This alteration depletes the pool of insulin-responsive vesicles marked by TUG and Usp25m. Mistargeting of GLUT4 and IRAP may contribute to distinct aspects of the metabolic syndrome in humans.

---

Insulin resistance is a core pathophysiologic component of type 2 diabetes and is associated with cardiovascular disease, hypertension, and obesity^1–3^. Reduced insulin action results from impairment of proximal steps in insulin signaling, yet data also implicate alteration of a distal step in the regulation of glucose uptake^1,3,4^. To stimulate glucose uptake, insulin acts in muscle and fat cells to mobilize intracellular vesicles containing GLUT4 glucose transporters^5^. In humans with insulin resistance, GLUT4 is mislocalized during fasting, so that it is depleted from insulin-responsive intracellular membranes^6–8^. These data, obtained ∼30 years ago, suggested that insulin resistance results in part from mistargeting of GLUT4, independent of an insulin signal. Yet, because GLUT4 regulation was not understood, studies to test this hypothesis could not be performed until now.

The main, insulin-regulated intracellular compartment containing GLUT4 is the pool of GLUT4 Storage Vesicles (GSVs), also called Insulin-Responsive Vesicles^5^. These vesicles accumulate during fasting and bind peripheral membrane proteins to enable insulin action^9–12^. Specifically, at the endoplasmic reticulum (ER)-Golgi intermediate compartment (ERGIC), GLUT4 and the transmembrane aminopeptidase IRAP are co-recruited into GSVs during vesicle assembly.

These proteins engage regulatory proteins including TUG (also called Aspscr1 or UBXN9), AS160 (also called Tbc1d4), and Usp25m, a splice form of the Usp25 protease that is present in adipose and muscle^10,13,14^. TUG, AS160, and Usp25m are each required for insulin-regulated GLUT4 translocation in adipocytes, and AS160 and TUG are required in muscle^13,15–25^.

The TUG protein traps GSVs intracellularly^5,14,16,19^. The N-terminal region of TUG binds GLUT4 and IRAP^19,26^, and central and C-terminal regions bind the ERGIC and Golgi matrix^27–30^. Insulin triggers site-specific cleavage of TUG to liberate the GSVs, mobilizing GLUT4^13,15,16,19,28^. In adipocytes, cleavage is mediated by Usp25m and divides intact TUG into 18 kDa N-terminal and 42 kDa C-terminal products^13^. In muscle, TUG deletion results in 85-90% of the effect of insulin to translocate GLUT4, which increases glucose uptake during fasting^15,16,31^. After TUG is cleaved, its C-terminal product regulates transcription to enhance thermogenesis^16^.

In rodents with high-fat diet (HFD)-induced insulin resistance, the Usp25m protease is depleted and TUG cleavage is attenuated^13,16^. Conversely, in humans, caloric restriction potentiates insulin action, increases Usp25m abundance, and decreases TUG abundance in adipose^32^. In HFD-fed mice, the reduction in adipose Usp25m abundance is post-transcriptional^13^. Possibly, Usp25m is destabilized. We hypothesize that in HFD-induced muscle insulin resistance, the establishment of a pool of GSVs and their regulatory proteins is reduced, so that fewer GSVs accumulate during fasting.

Here, we show first that Usp25m is required for GLUT4 translocation and glucose uptake in muscle. The phenotype of muscle-specific *Usp25* knockout (MUKO) mice is ablated by HFD feeding, concurrent with Usp25m depletion in control mice. Moreover, GSV-associated proteins act together: AS160 is required for efficient TUG cleavage, and Usp25m is required for co-trafficking of IRAP and GLUT4. HFD feeding also dissociates IRAP and GLUT4 co-targeting, and ablates the previously-shown effect of muscle *Tug* (*Aspscr1*) knockout (MTKO) to cause increased glucose uptake during fasting. We conclude that HFD feeding depletes the pool of TUG- and Usp25m-regulated GSVs. Our data identify a new locus of insulin resistance, characterized by insulin-independent mislocalization of GLUT4 and IRAP.

## Results

### Usp25m deletion impairs insulin-stimulated glucose uptake in muscle

Previous data show that the Usp25m protease is required for TUG cleavage and GLUT4 translocation in 3T3-L1 adipocytes^13^. To test if Usp25m is similarly required in muscle and to study its role in glucose homeostasis, we created muscle Usp25 knockout (MUKO) mice. As shown in Fig. 1a, MUKO mice had deletion of Usp25 in skeletal muscle, but not in other tissues. We treated these mice with intraperitoneal (IP) injection of insulin and glucose, or saline control, then euthanized them 30 min. after IP injection. In quadriceps of WT control mice, insulin caused the generation of a 42 kDa TUG C-terminal cleavage product, similar to previous results^13,16^, but in MUKO quadriceps, TUG cleavage was attenuated (Fig. 1b). These results support the idea that Usp25m is required for insulin-stimulated endoproteolytic TUG cleavage in skeletal muscle.

**Fig. 1.**
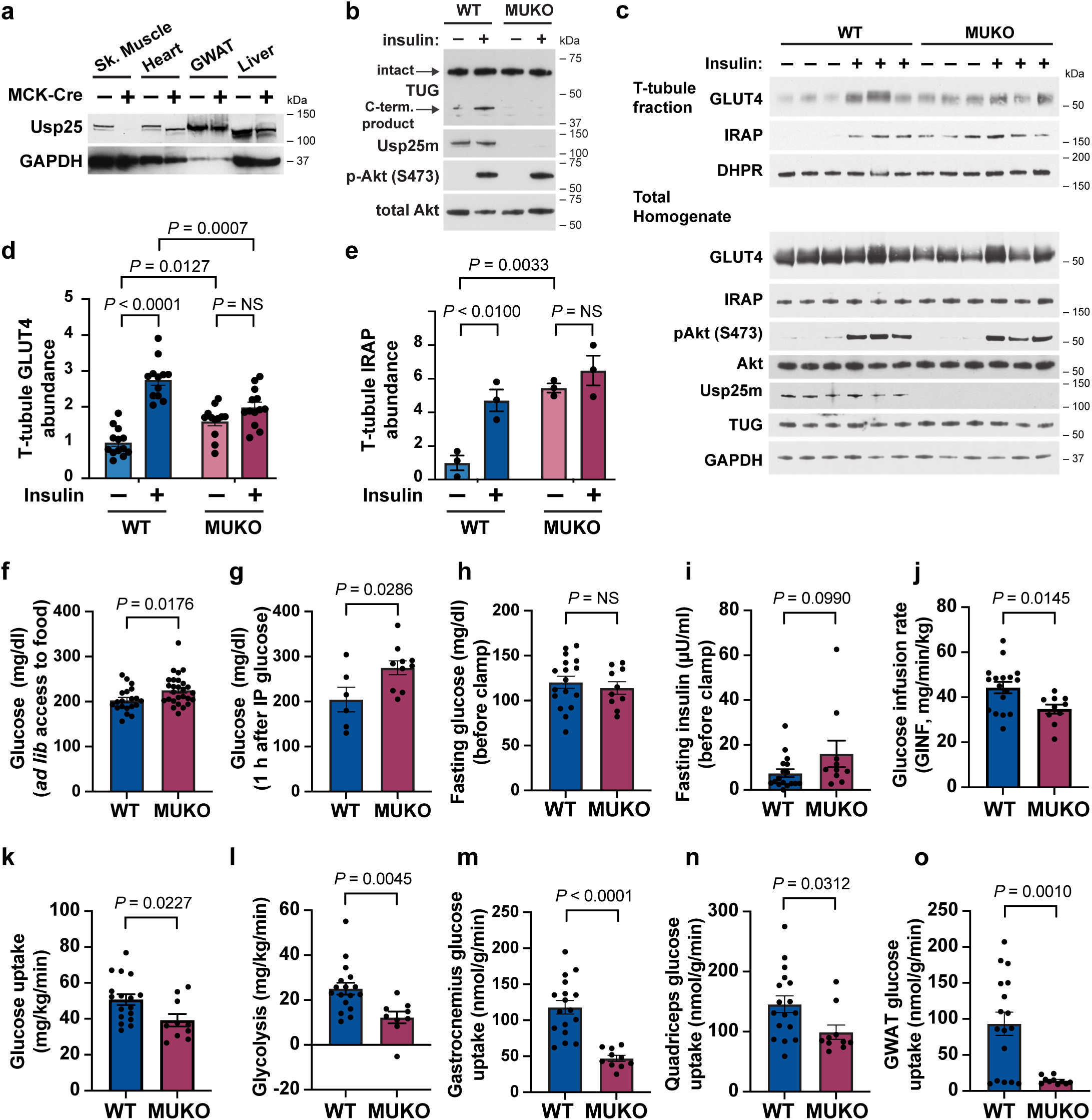
Mice with Usp25 deletion in muscle have impaired insulin-stimulated glucose uptake. **a.** Immunoblotting was performed as indicated on tissues from MUKO and WT control mice. Sk. Muscle, hindlimb skeletal muscle; GWAT, gonadal white adipose tissue. **b.** MUKO and WT control mice were fasted 6 h, then treated with IP injection of insulin and glucose, or saline control, then euthanized after 30 min. Quadriceps muscles were isolated and immunoblotted as indicated. **c.** MUKO and WT control mice were treated as in (b). Quadriceps muscles were isolated and homogenized, and T-tubule enriched membrane fractions were isolated. Immunoblots were done as indicated. **d. ,e.** Data from mice treated as in (c) were quantified. Abundances of GLUT4 (d) and IRAP (e) are shown. For GLUT4 (d), n = 13 for WT basal, 12 for WT insulin-stimulated, 12 for MUKO basal, and 13 for MUKO insulin-stimulated. For IRAP (e), n = 3 in each group. NS, not significant. **f.,g.** Plasma glucose was measured in 12-week-old MUKO and WT control mice with *ad lib* access to food (f) and after a glucose load (g). Mice in (g) were fasted for 6 h, then treated IP with 2 g/kg glucose, and measurements were made 1 h after IP injection. In (f), n = 20 WT and 27 MUKO mice. In (g), n = 6 WT and 10 MUKO mice. **h.-o.** Hyperinsulinemic-euglycemic clamps were performed in 12-week-old MUKO and WT control mice. Plots show fasting glucose and insulin concentrations prior to the clamp (h,i), glucose infusion rate (GINF) during steady state (6 time points at 90-140 min. of the clamp; j), whole body glucose uptake and glycolysis (k,l), and tissue-specific glucose uptake in gastrocnemius (m), quadriceps (n), and GWAT (o). n = 17 WT and 10 MUKO mice. All data are presented as mean ± s.e.m. of biologically independent samples, analyzed using a two-tailed t test or ANOVA with adjustment for multiple comparisons.

To test the prediction that Usp25m is required for GLUT4 translocation, we prepared T-tubule membrane fractions from quadriceps muscles. Wildtype (WT) and MUKO mice were treated with IP insulin and glucose, or saline control, then mice were euthanized and quadriceps muscles were isolated. GLUT4 and IRAP abundances were increased by insulin treatment in T-tubules prepared from WT mice, and this effect of insulin was abrogated in MUKO mice (Fig. 1c and Suppl. Fig. 1a). Quantification of data from several mice show that insulin caused a 2.8-fold increase in T-tubule GLUT4 abundance in wildtype mice (Fig. 1d). In MUKO mice, GLUT4 was increased 1.6-fold in T-tubules from unstimulated muscles, and there was no further increase in insulin-stimulated muscles. Insulin-stimulated translocation of IRAP was also disrupted by Usp25m deletion, as predicted (Fig. 1c). Insulin stimulated a 4.7-fold increase in IRAP abundance in T-tubules of WT muscles (Fig. 1e). In T-tubules from MUKO mice, T-tubule IRAP abundance was similarly increased (by 5.5-fold) in the absence of insulin stimulation, and there was no further effect of insulin (Fig. 1e). Control immunoblots of total homogenates showed that Usp25 knockout caused a 25% reduction in GLUT4 abundance; no effects on IRAP abundance or Akt phosphorylation were observed (Fig. 1c, Suppl. Fig. 1b). Differential effects of Usp25m knockout on GLUT4 and IRAP targeting were quantified, and reveal that the ratio of IRAP to GLUT4 abundances in T-tubules was increased in MUKO animals, compared to WT controls (Suppl. Fig. 1c,d). This increase was 2.6-fold in unstimulated muscles, and 2.4-fold in insulin-stimulated muscles. This dissociation of IRAP and GLUT4 targeting suggests that Usp25m deletion may not only impair the release of GSVs to the cell surface, but may also impair the co-recruitment of IRAP and GLUT4 into GSVs, as discussed below. The main point is that Usp25m is required for insulin-responsive trafficking of GLUT4 and IRAP in muscle.

We next studied effects on glucose homeostasis. MUKO mice with *ad libitum* access to chow had increased blood glucose concentrations, compared to WT controls (Fig. 1f). Blood glucose concentrations were unchanged in 6 h fasted MUKO mice, but were increased compared to those in control mice after IP glucose injection (Fig. 1g, Suppl. Fig. 1e). For hyperinsulinemic-euglycemic clamp studies, MUKO and WT control mice were fasted overnight. Baseline studies revealed no differences in fasting plasma glucose concentrations or in body weight or composition (Fig. 1h, Suppl. Fig. 1f-h). Plasma insulin concentrations trended toward an increase (by 2.2-fold) in MUKO mice, compared to controls (Fig. 1i). There was a trend toward increased HOMA-IR, a measure of insulin resistance during fasting (Suppl. Fig. 1i). In clamp studies, the glucose infusion rate required to maintain euglycemia was reduced by 22% in MUKO mice, compared to controls (Fig. 1j, Suppl. Fig. 1j-m). This was due to a 23% decrease in whole-body glucose uptake in MUKO mice (Fig. 1k). Glycolysis was decreased by 51% (Fig. 1l). There were no genotype-dependent alterations in endogenous glucose production under fasting or hyperinsulinemic-euglycemic conditions (Suppl. Fig. 1n,o). In MUKO mice, compared to controls, gastrocnemius muscle glucose uptake was reduced by 60%, and quadriceps muscle glucose uptake was reduced by 32% (Fig. 1m,n). No change in glucose uptake was observed in the heart, which did not have deletion of Usp25m (Suppl. Fig. 1p and Fig. 1a). We observed an 84% reduction in glucose uptake in gonadal white adipose tissue (GWAT) in MUKO mice, compared to controls, which may reflect chronic hyperinsulinemia (Fig. 1o). We noted enhanced suppression of circulating nonesterified fatty acids (NEFA) during the clamp in MUKO mice, compared to controls (Suppl. Fig. 1q-s). The data show that muscle-specific deletion of Usp25m causes a dramatic reduction in insulin-stimulated glucose uptake, observable in both whole-body analyses and in individual skeletal muscles.

### MUKO mice have reduced energy expenditure normalized to lean body mass

The TUG C-terminal cleavage product promotes thermogenesis^16^. Thus, we considered that MUKO mice might have reduced energy expenditure, resulting from impaired TUG cleavage. Consistent with this idea, indirect calorimetry showed that energy expenditure was reduced by 6% in MUKO mice, compared to WT controls, when normalized to lean mass (Fig. 2a,b). This effect was more marked during dark hours than during light hours, and was observed on a per mouse basis across a range of lean masses (Fig. 2c-e). The rate of oxygen consumption (VO_2_) was decreased in MUKO mice, compared to controls, yet no changes in the rate of carbon dioxide production (VCO_2_) were detected (Fig. 2f,g). The respiratory exchange ratio (RER) was not significantly changed (Fig. 2h). When normalized to total body weight, there remained a trend toward reduced energy expenditure in MUKO mice, which was not significant (Suppl. Fig. 2a-e). No genotype-dependent changes were observed in locomotor activity, food intake, or water intake (Fig. 2i,j, Suppl. Fig. 2f-h). Total body weight, fat mass, lean mass, and percent fat mass were unchanged in MUKO mice, relative to controls (Fig. 2k-m, Suppl. Fig. 2i). There was a slight (1.3%) increase in percent lean mass, which cannot account for the decreased energy expenditure we observed (Suppl. Fig. 2j). Together, the data show that MUKO mice have reduced energy expenditure, compared to WT controls. The data support the idea that impaired TUG cleavage causes impairments in both glucose uptake and energy expenditure.

**Fig. 2.**
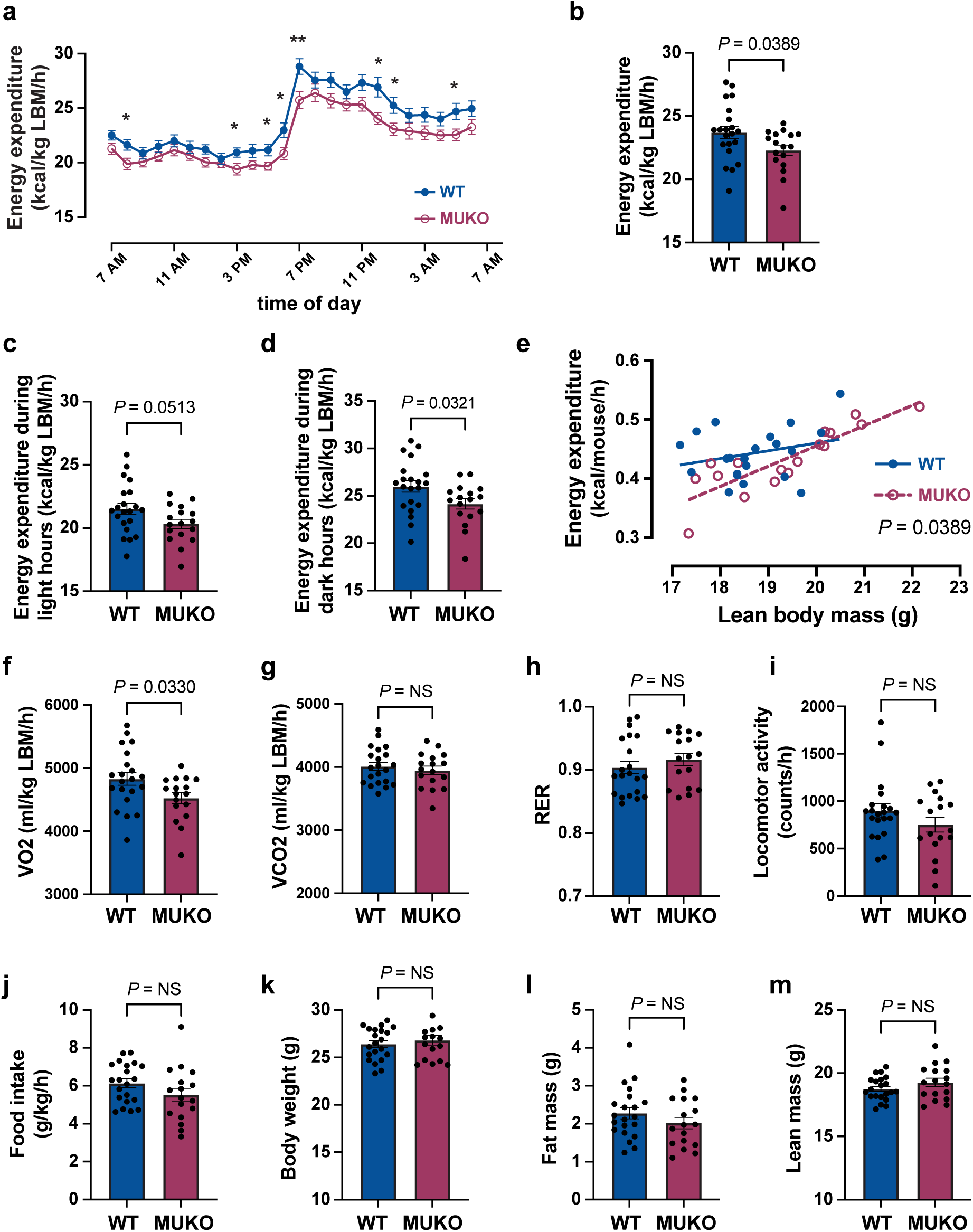
Mice with Usp25m deletion in muscle have reduced energy expenditure on regular chow. **a.** Energy expenditure was measured by indirect calorimetry in 13-week-old mice and is plotted over time. n = 21 WT and 17 MUKO mice. Mean ± s.e.m. is shown. Individual time points were analyzed using two-tailed t tests. \**P*<0.05, \*\**P*<0.01. **b.-d.** Energy expenditure in (a) was averaged over 24 h, during light hours (c), and during dark hours (d). n = 21 WT and 17 MUKO mice. LBM, lean body mass. **e.** Energy expenditure per mouse is plotted as a linear regression versus lean body mass. n = 21 WT and 17 MUKO mice. **f.-m.** The indicated parameters were measured in metabolic cages. n = 21 WT and 17 MUKO mice. All data are presented as mean ± s.e.m. of biologically independent samples, analyzed using a two-tailed t test (b-d,f-m) or ANCOVA (e).

### High-fat diet feeding ablates phenotypes observed in MUKO mice

In mice with diet-induced insulin resistance, impaired insulin signaling may reduce TUG cleavage. Yet, previous data show that high-fat diet (HFD) feeding causes a marked reduction of Usp25m protein abundance in both adipose and muscle^13,16^. We predicted that the phenotypes we observed in MUKO mice might be reduced or ablated in mice fed a HFD, at least in part because of reduced Usp25m abundance in control mice. Supporting this idea, we observed no genotype-dependent differences in hyperinsulinemic-euglycemic clamp studies performed on mice fed a HFD for three weeks. After the HFD, MUKO and control mice had similar body weight and composition (Suppl. Fig. 3a-e). Fasting glucose and insulin concentrations (before the clamp), glucose infusion rates, fasting and clamped endogenous glucose production, and glycolysis were all unchanged in MUKO mice (Fig. 3a-d, Suppl. Fig. 3f-k). Whole-body and tissue-specific glucose uptake were also similar in HFD-fed MUKO and WT control mice (Fig. 3e-h, Suppl. Fig. 3l).

**Fig. 3.**
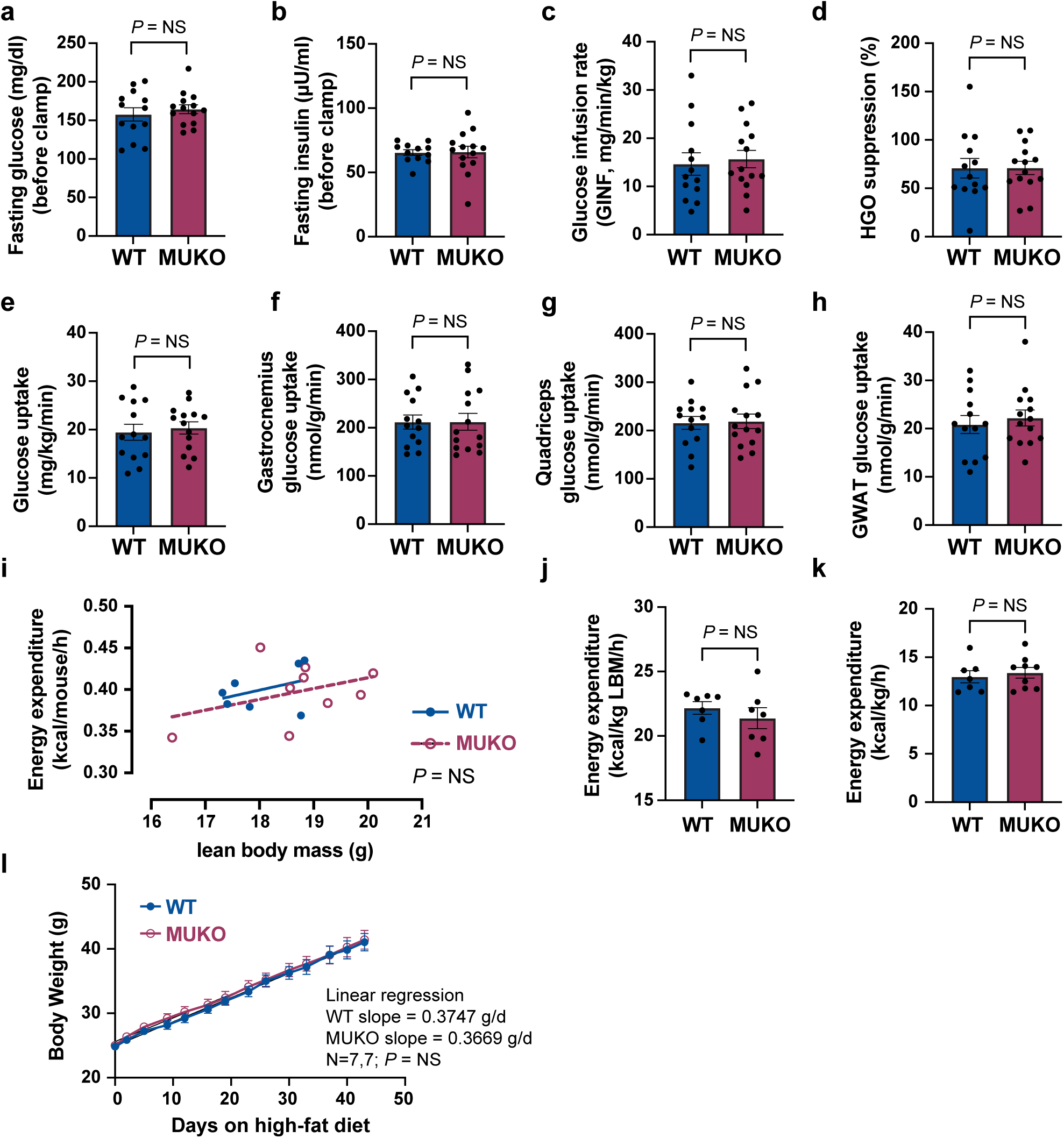
High-fat diet feeding ablates the effects of muscle Usp25m knockout on glucose uptake and energy expenditure. **a.-h.** Mice were treated with a high-fat diet (HFD) for 3 weeks, then studied in hyperinsulinemic-euglycemic clamps at age 15 weeks. Basal glucose and insulin concentrations were measured prior to the clamps (a,b), and the indicated parameters were measured during clamps and are plotted. n = 13 WT and 14 MUKO mice. Mean ± s.e.m. is shown, analyzed using two-tailed t tests. GINF, glucose infusion rate. EGO, endogenous glucose production. GWAT, gonadal white adipose tissue. NS, not significant. **i.-k.** Mice were fed a HFD for 3 weeks and studied in metabolic cages at age 12 weeks. Plots show energy expenditure per mouse as a linear regression versus lean body mass (i), energy expenditure per lean body mass (LBM, j), and energy expenditure per total body weight (k). n = 7 WT and 9 MUKO mice. Data in (i) were analyzed by ANCOVA. Data in (j,k) are shown as mean ± s.e.m., analyzed using a two-tailed t test. NS, not significant. **l.** Mice were fed a HFD beginning at 8 weeks of age, and body weights were measured and are plotted versus number of days on the HFD. A linear regression was used to assess the rate of weight gain, and was not significantly different in MUKO mice, compared to WT controls. n = 7 in each group, shown as mean ± s.e.m.

Compared to RC-fed mice studied above, the HFD-fed mice had markedly reduced rates of glucose infusion and whole-body glucose uptake, indicating that the HFD did indeed caused insulin resistance. Circulating NEFA concentrations were also unchanged in HFD-fed MUKO mice, compared to controls (Suppl. Fig. 3m-o). The data show that the effects of muscle Usp25m deletion to impair insulin-stimulated glucose uptake in RC-fed mice are no longer observed when mice are fed a HFD.

We also observed no difference in energy expenditure in HFD-fed MUKO mice, compared to controls. In these experiments, we used indirect calorimetry (Fig. 3i-k, Suppl. Fig. 4a-i). There were no genotype-dependent differences in energy expenditure (normalized to lean or total body mass), VO_2_ or VCO_2_, RER, locomotor activity, or food or water intake. Compared to RC-fed controls, above, HFD-fed mice had a slight reduction in energy expenditure. HFD-fed MUKO mice had a slight (4%) increase in lean mass, with no difference in fat mass (Suppl. Fig. 4i,h).

When body weight was followed over time after beginning a HFD, there was no genotype-dependent difference in the rate of weight gain, final body weight or GWAT mass (Fig. 3l, Suppl. Fig. 4j,k). Thus, the reduced energy expenditure observed in MUKO mice fed a RC diet is no longer observed when mice are fed a HFD.

### GSV-regulating proteins act together and this action is disrupted in HFD-fed mice

Previous data support the idea that transmembrane (GLUT4, IRAP) and non-transmembrane proteins (TUG, AS160, Usp25m) proteins interact to regulate GSVs in fat and muscle cells^14^. AS160, TUG, and tankyrase (TNKS1 and TNKS2, collectively TNKS) bind to adjacent peptides within the cytosolic N-terminus of IRAP, and TUG and TNKS recruit Usp25m, as depicted in Fig. 4a. Endogenous AS160 and Usp25m reside on TUG-bound GSVs in 3T3-L1 adipocytes^13,21,33^. We observed that TUG coimmunoprecipitates with AS160 and with TNKS2, and the ankryrin repeat domain of TNKS2 was sufficient for this interaction (Suppl. Fig. 5a,b). Previous data also show that co-expression of AS160 markedly enhances the effect of Usp25m to cleave TUG in transfected cells^13^.

**Fig. 4.**
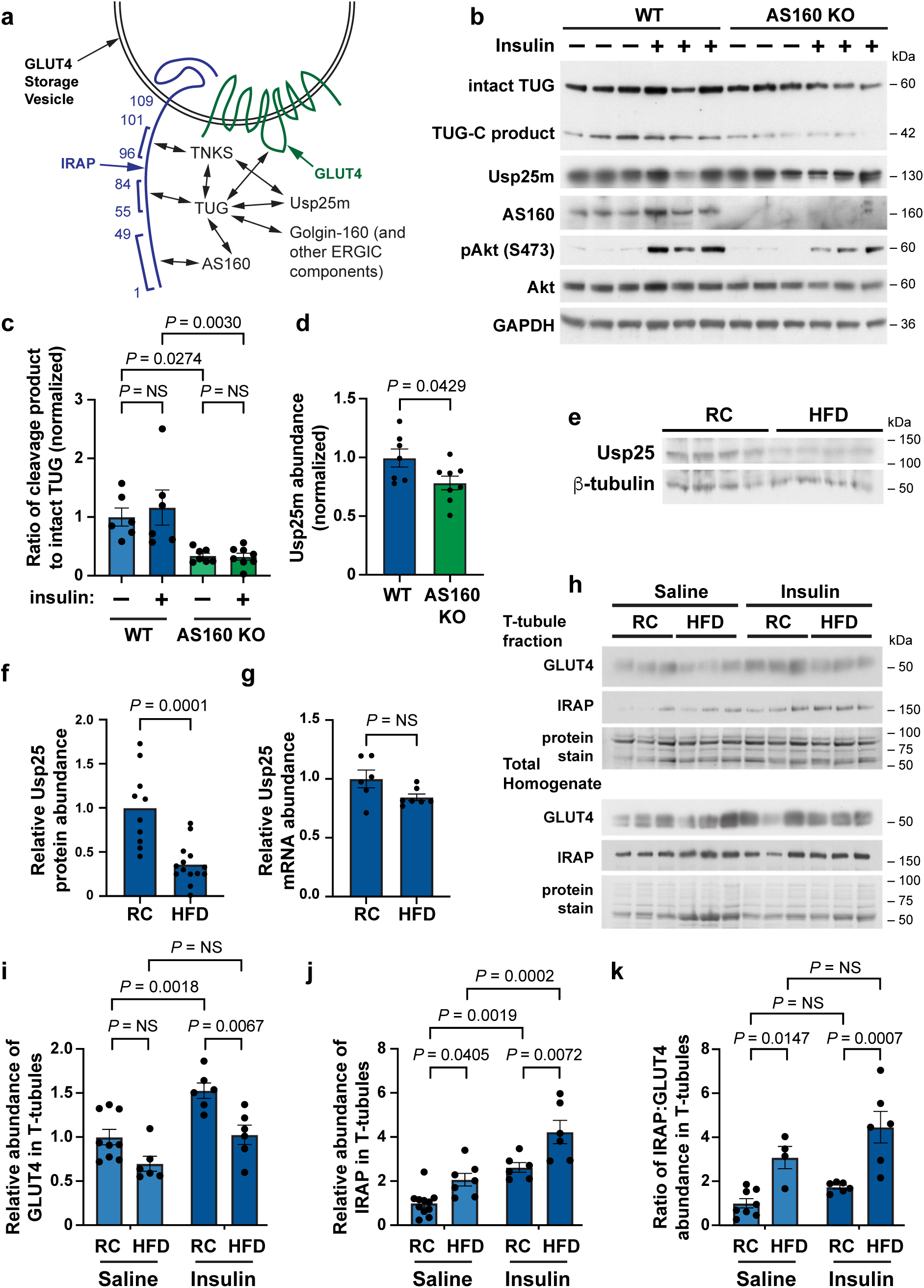
Proteins that regulate GLUT4 storage vesicles act together and this action is disrupted in HFD-fed mice. **a.** A diagram of interactions among proteins that regulate GLUT4 Storage Vesicles. Residues in the cytosolic N-terminus of IRAP are numbered, and AS160, TUG, and TNKS bind the indicated peptides. Interactions among proteins are indicated by arrows. The complex is anchored to Golgin-160 and other components at the endoplasmic reticulum-Golgi intermediate compartment (ERGIC). Modified from Ref. ^14^. **b.-d.** Epitrochlearis muscles were isolated from WT and AS160 knockout (KO) rats, treated with or without insulin, then lysates were prepared and immunoblotted (b). Intact TUG and its C-terminal cleavage product (TUG-C product) are indicated. The ratio of the cleavage product to intact TUG abundance was quantified (c). Usp25m abundance in unstimulated muscles was quantified (d). Data were normalized to that in WT controls. n = 6 WT unstimulated, 6 WT insulin-stimulated, 7 KO unstimulated, and 8 KO insulin-stimulated samples. **e.-g.** Mice were fed regular chow (RC) or high-fat diet (HFD) for 3 weeks until age 13 weeks, then hindlimb muscles were isolated and immunoblotted as indicated (e). Usp25m protein abundance was quantified, normalized to β-tubulin abundance, and plotted (f). n = 10 RC and 14 HFD samples. In parallel, Usp25m mRNA abundance was quantified and is plotted (g). 6 WT and 7 HFD samples. **h.-k.** Mice were fed RC or HFD for 6 weeks, beginning at 8 weeks of age. Mice (14 weeks old) were then fasted 6 h and treated with IP injection of insulin-glucose solution, or saline control. After 30 min., mice were euthanized, quadriceps muscles were homogenized, and T-tubule membrane fractions were isolated. Immunoblots were done as indicated (h). The relative abundances of GLUT4 and IRAP in T-tubule membranes as quantified and is plotted (i,j). For GLUT4 (i), n = 9 RC saline, 6 RC insulin, 6 HFD saline, and 6 HFD insulin samples. For IRAP (j), n = 11 RC saline, 7 HFD saline, 6 RC insulin, and 6 HFD insulin samples. The ratio of IRAP to GLUT4 abundances was quantified and is plotted (k). n = 8 RC saline, 4 HFD saline, 6 RC insulin, and 6 HFD insulin samples. All data are presented as mean ± s.e.m. of biologically independent samples, analyzed using a two-tailed t test (d,f,g) or ANOVA (c,I,j,k). NS, not significant.

To test whether AS160 is required for insulin-stimulated TUG cleavage, we used isolated epitrochlearis muscles from AS160 KO and WT control rats^34,35^. Muscles were treated with or without insulin, lysed and immunoblotted to detect intact TUG and its C-terminal cleavage product, Usp25m, and control proteins. As shown in Figs. 4b,c, the abundance of the TUG C-terminal product, relative to intact TUG, was reduced by ∼72% in insulin-treated AS160 KO versus control muscles. Reduced TUG cleavage product was also observed in unstimulated AS160 KO muscles. There was no marked effect of ex vivo insulin stimulation to increase the abundance of the cleavage product in WT muscles, possibly because only cells at the surface of the muscle fiber are stimulated under these conditions. The overall abundance of intact TUG protein was not changed in unstimulated AS160 KO muscles, compared to WT controls (Suppl. Fig. 5c). We observed a ∼21% decrease in the abundance of Usp25m in unstimulated AS160 KO muscles (Fig. 4d). We conclude that AS160 is required for efficient TUG cleavage in muscle.

Previous data show that Usp25m protein abundance is reduced in insulin resistance and that, in adipocytes, this occurs without any change in its mRNA abundance^13,16^. To test if HFD-induced effects in muscle are similarly post-transcriptional, we measured Usp25m protein and mRNA abundances in hindlimb muscles. Usp25m protein abundance was reduced by ∼65% in muscles from HFD-fed mice, compared to RC-fed controls, similar to previous results (Fig. 4e,f). In the same tissues, Usp25 mRNA abundance was unchanged (Fig. 4g). These data show that the effect of HFD feeding to cause decreased Usp25m protein abundance in muscle is post-transcriptional. The data suggest that Usp25m may be destabilized in insulin resistant mice.

Because destabilization of proteins often occurs when protein complexes are not properly assembled, we considered that a complex containing Usp25m, similar to that diagrammed in Fig. 4a, might not be well-formed in HFD-induced insulin resistance. To test if membrane proteins in this complex are mislocalized in HFD-fed mice, we isolated T-tubules from quadriceps muscles and immunoblotted IRAP and GLUT4. In HFD-fed mice, compared to RC-fed controls, the abundance of GLUT4 in T-tubules was not altered during fasting; as expected, the effect of insulin to increase in T-tubule GLUT4 was impaired (Fig. 4h,i). In contrast, T-tubule IRAP abundance was increased in HFD-fed mice, compared to RC-fed controls, both during fasting and after insulin stimulation (Fig. 4h,j). Overall, there was an approximately twofold increase in IRAP in T-tubules from HFD-fed mice, compared to RC-fed controls (Suppl. Fig. 5d). As well, the ratio of IRAP to GLUT4 in T-tubule membranes was increased in HFD-fed mice, both during fasting and after insulin stimulation (Fig. 4k). Overall, the ratio of IRAP to GLUT4 in T-tubule membranes was increased threefold in HFD-fed mice, compared to RC-fed controls (Suppl. Fig. 5e). Because IRAP and GLUT4 are normally co-recruited into GSVs during vesicle formation, and trapped intracellularly in these vesicles, the increased ratio of IRAP to GLUT4 in T-tubules of fasting mice suggests that the pool of GSVs is depleted in HFD-fed mice.

To further test the idea that an increased ratio of IRAP to GLUT4 in T-tubules can reveal an impairment in GSV formation, we studied muscle TUG knockout (MTKO) mice. Previous data show that GSVs are formed normally in 3T3-L1 adipocytes with shRNA-mediated depletion of TUG^20^. Specifically, vesicles characteristic of GSVs, and distinct from endosomes, are observed fusing directly at the plasma membrane in these cells in the absence of insulin stimulation. In MTKO mice, the increases in IRAP and GLUT4 in T-tubule membranes of fasted mice are similar to those observed after insulin stimulation of control mice^16^. We further quantified these data, and observed that insulin stimulation and TUG deletion both cause a decrease in the ratio of IRAP to GLUT4 in T-tubules (Suppl. Fig. 5f,g). No further effect of insulin was observed in MTKO mice. Thus, the effects of TUG deletion or insulin stimulation on the T-tubule IRAP to GLUT4 ratio are opposite to those observed in HFD-induced insulin resistance (Fig. 4k, Suppl. Fig. 5e). The data support the idea that, in RC-fed mice, IRAP and GLUT4 are co-recruited into TUG-bound GSVs, and are thus trapped intracellularly. In HFD-induced insulin resistance, impaired establishment of the pool of GSVs disrupts this entrapment, and results in an increased ratio of IRAP to GLUT4 in T-tubule membranes of fasting mice.

### A high-fat diet ablates the effect of muscle TUG knockout to increase glucose uptake during fasting

To test definitively whether TUG-bound GSVs are depleted in HFD-induced insulin resistance, we studied MTKO mice during fasting. In contrast to Usp25m deletion, which causes impaired GSV release, TUG deletion causes constitutive mobilization of the GSVs, similar to effects of insulin stimulation. This effect occurs during fasting, making it possible to study effects on the GSVs themselves, in the absence of an insulin signal. Previous data show that, on a RC diet, fasting MTKO mice have reduced glucose and insulin concentrations, a 27% increase in whole-body glucose uptake, and 2.0-and 1.8-fold increases in tissue-specific glucose uptake in gastrocnemius and quadriceps muscles, respectively^16^. We reasoned that if the pool of TUG-regulated GSVs is ablated by HFD feeding, then these effects of TUG deletion would be lost in HFD-fed mice. Accordingly, we treated MTKO and WT control mice with a HFD and tested whether this ablated the effects of TUG deletion on fasting glucose homeostasis.

To induce insulin resistance without causing genotype-dependent changes in body weight or composition, we studied younger mice than previously^16^. Mice were placed on a HFD beginning at age 8 weeks, indwelling jugular catheters were placed during the third week of the diet, and glucose turnover studies were done at age 12 weeks (see methods). As shown in Fig. 5a and 5b, there were no differences in fasting plasma insulin or glucose concentrations in HFD-fed MTKO mice, compared to WT controls. Accordingly, HOMA-IR was not different between MTKO and WT mice (Suppl. Fig. 6a), yet confirmed that mice were insulin resistant compared to similarly aged RC-fed mice (Suppl. Fig. 1i). No differences were observed in body weight, fat or lean mass, or percent fat or lean mass (Suppl. Fig. 6b-f).

**Fig. 5.**
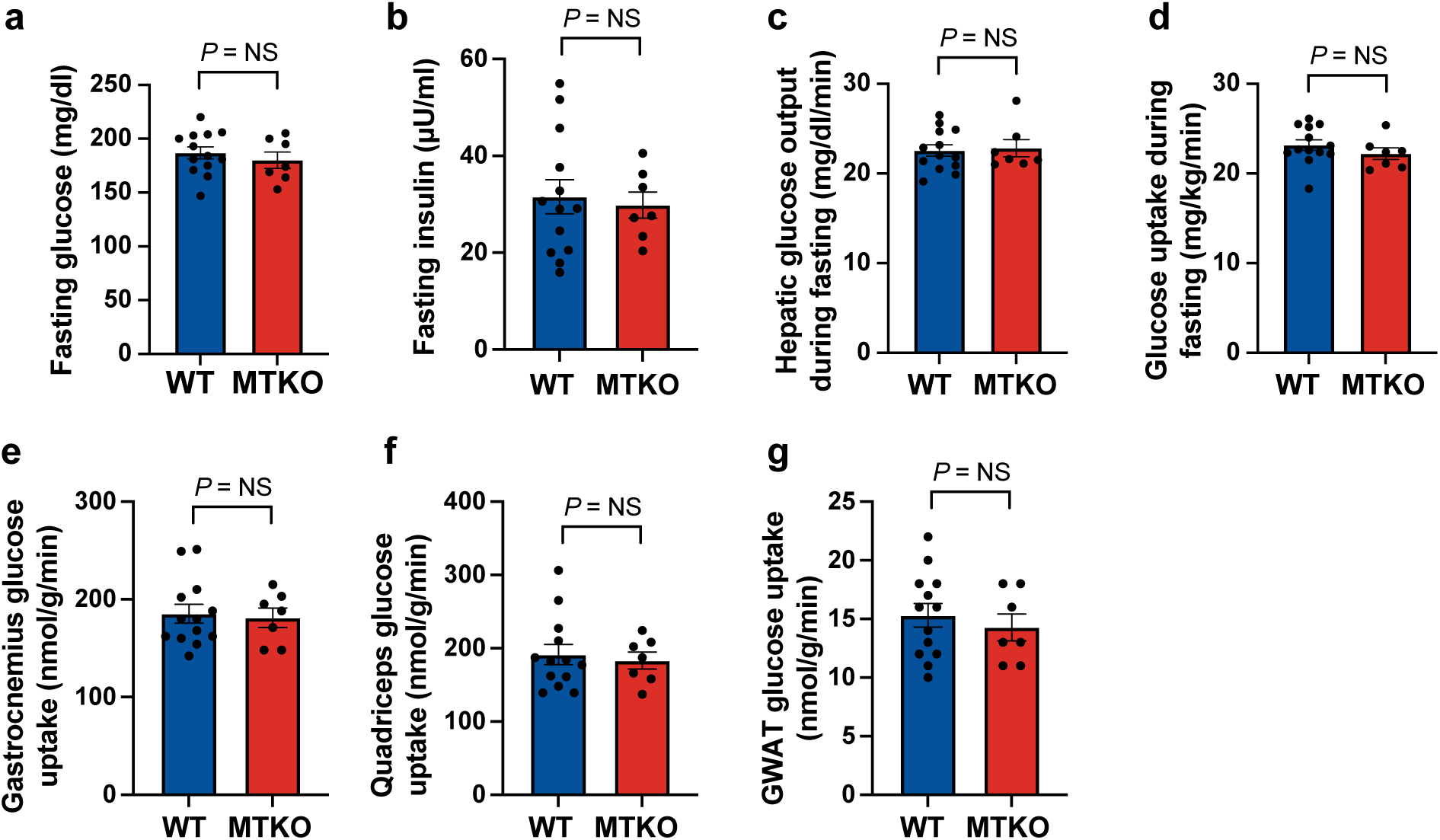
Effects of muscle-specific TUG knockout to increase fasting glucose uptake are not observed in high-fat diet -fed mice. **a.-g.** Muscle TUG knockout (MTKO) and WT control mice were previously studied and, on a RC-diet, MTKO mice had increased whole-body and muscle-specific glucose uptake. Here, MTKO and WT control mice were fed a HFD starting at 8 weeks of age, and glucose turnover studies were performed, as previously in fasting mice, after 3.5 weeks on the HFD. Basal glucose and insulin concentrations were measured prior to the study (a,b), and the indicated parameters were measured during the study (c-g) and are plotted. n = 13 WT and 7 MTKO mice. Data are presented as mean ±s.e.m., and analyzed using a two-tailed t test. GWAT, gonadal white adipose tissue. NS, not significant.

Dynamic measurements of glucose flux were assessed in fasting MTKO and WT control mice using tracers, as previously^16^. We observed no differences in endogenous glucose production or whole-body glucose turnover (Fig. 5c,d). Plasma glucose concentrations were similar in MTKO and WT control mice during these studies (Suppl. Fig. 6g). No differences were observed in insulin concentrations at the end of the infusion, in rate of glycolysis, or in non-esterified fatty acid concentrations (Suppl. Fig. 6h-j). We measured tissue-specific glucose uptake using a 2-deoxyglucose tracer, and observed no genotype-dependent differences in gastrocnemius or quadriceps muscle-specific glucose uptake (Fig. 5e,f). As well, no differences were observed in tissue-specific glucose uptake in GWAT or heart (Fig. 5g; Suppl. Fig. 6k). These results are markedly different from those obtained in the same mice, using the same experimental protocol, on a RC diet^16^. Together, the data show that HFD-induced insulin resistance is characterized by alterations in GLUT4 targeting during fasting, so that the effect of TUG to trap GLUT4 in an intracellular, insulin-responsive pool of GSVs is no longer observed.

## Discussion

Our data show that the Usp25m protease is required for full effects of insulin to stimulate TUG cleavage, GLUT4 translocation and glucose uptake in skeletal muscle. Deletion of Usp25m in muscle also caused reduced energy expenditure, normalized to lean mass, consistent with loss of the effect of the TUG C-terminal cleavage product to enhance thermogenesis. In HFD-fed mice, Usp25m abundance was reduced post-transcriptionally and TUG cleavage was impaired, so that these effects of Usp25m knockout were no longer observed. The data suggest that the establishment of a GSV pool and the formation of a GSV-regulating protein complex are interdependent processes, which are lost in HFD-induced insulin resistance. Further supporting this idea, effects of muscle-specific TUG knockout to increase glucose uptake during fasting were ablated in HFD-fed mice. The results support a model in which vesicle trafficking at this Usp25m-TUG locus is disrupted during fasting to contribute to insulin resistance in muscle (Fig. 6). More broadly, the size of the pool of GSVs may be a key determinant of insulin sensitivity. Previous studies show that GLUT4 and IRAP are mislocalized during fasting in humans with insulin resistance^4,7,8^. Advances in understanding GLUT4 regulation have now enabled us to define the site of GLUT4 and IRAP mislocalization using molecular markers.

**Fig. 6.**
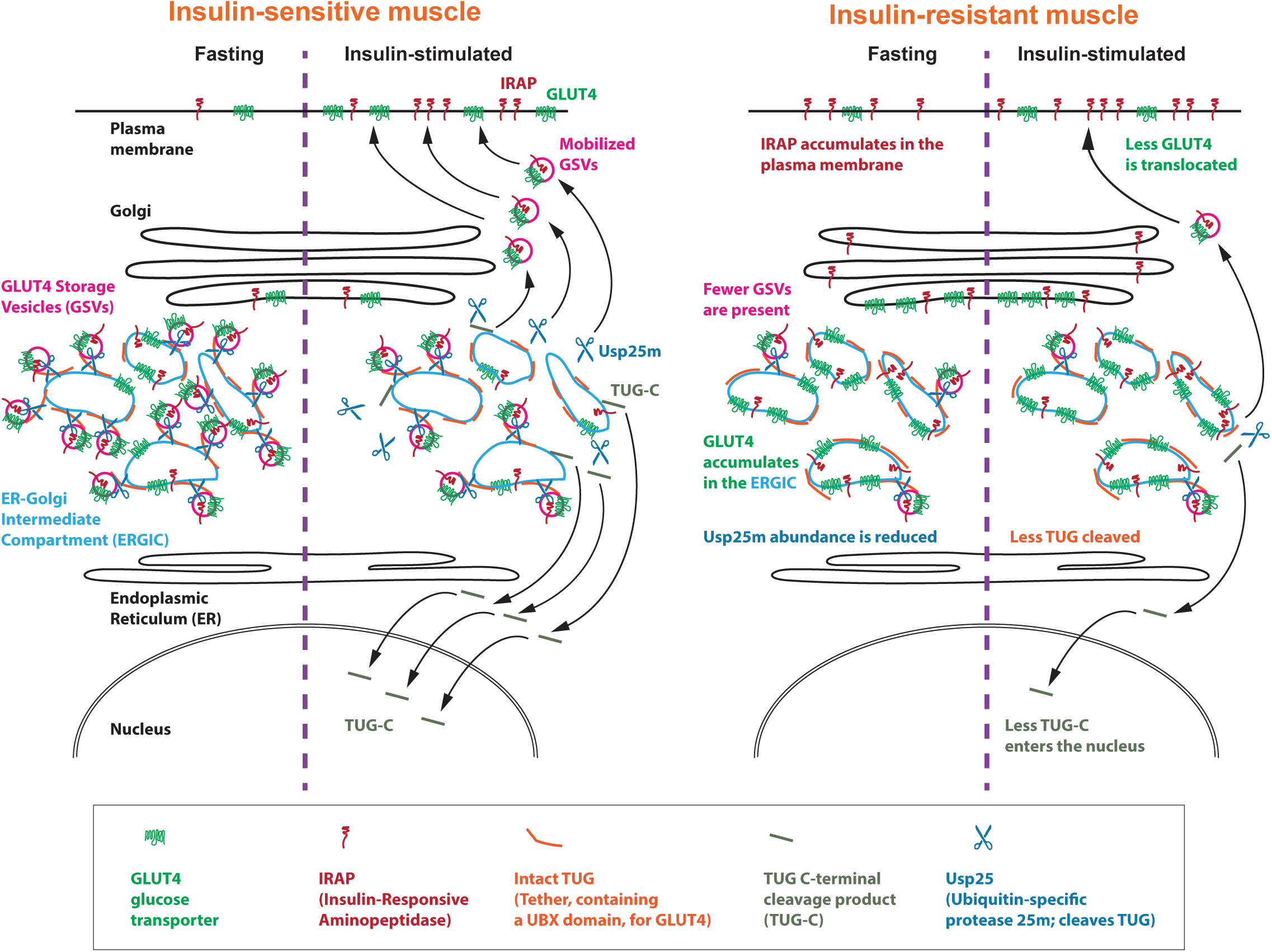
Model showing how insulin responsiveness is regulated by the size of a pool of insulin-responsive GLUT4 storage vesicles in muscle. In insulin-sensitive muscle (left), GLUT4 storage vesicles (GSVs) accumulate during fasting. GSVs reside near the endoplasmic reticulum (ER)-Golgi intermediate compartment (ERGIC), contain GLUT4 and IRAP, and are bound by TUG and Usp25m proteins. AS160 is also bound (not shown). Insulin stimulates Usp25m-mediated TUG cleavage to mobilize the GSVs, which inserts GLUT4 and IRAP at the plasma membrane to mediate glucose uptake and vasopressin inactivation, respectively. The TUG C-terminal cleavage product enters the nucleus and regulates transcription to control thermogenesis. In insulin-resistant muscle (right), fewer GSVs are present during fasting, so that GLUT4 and IRAP are not well sequestered together. GLUT4 likely accumulates in ERGIC membranes, and IRAP travels to the plasma membrane. Usp25m protease abundance is reduced. Upon insulin stimulation, less TUG is cleaved, fewer GSVs are mobilized to translocate GLUT4, and less TUG C-terminal product enters the nucleus.

Our data support the concept that the main insulin-responsive compartment, the GSVs, are vesicles that are trapped intracellularly by TUG and that are released by TUG cleavage^13,14,16^. Why has the nature of these vesicles been so elusive? One reason is that much previous work has relied upon cultured cell models, such as L6 myocytes and 3T3-L1 adipocytes, which may not adequately replicate cell type -specific mechanisms present *in vivo* in muscle and adipose cells^13,14,28^. Particular experimental manipulations may further disrupt these mechanisms^36^. As well, the TUG protein contains intrinsically disordered regions and ubiquitin-like domains, and it is likely susceptible to degradation during sample preparation^28,30^. TUG may participate in a biomolecular condensate, so that regulatory proteins recruited to GSVs bind each other though low-affinity interactions^14,30^. In aggregate, these interactions are sufficient to trap the vesicles in an insulin-responsive configuration. Yet, this biology complicates biochemical analyses, and genetic approaches were required to identify and characterize TUG as a critical regulator of GLUT4 trafficking^19,37^. Three different genetic models have been used to disrupt TUG action in skeletal muscle, and to show that it accounts for the bulk of GLUT4 translocation and glucose uptake in this tissue^15,16,38^. Our present data add that Usp25m deletion in muscle inhibits TUG cleavage and GLUT4 translocation, and impairs insulin-stimulated glucose uptake. The results support the view that, *in vivo*, other sites of GLUT4 regulation are relatively less important for the overall control of acute, insulin-responsive glucose uptake.

In muscle, insulin resistance comprises both a shift in the insulin dose response and a reduction in the maximal effect of insulin to stimulate glucose uptake^3,39^. The former effect may reflect an impairment of proximal steps in insulin signaling^2^. The latter effect may be more compatible with a reduction in the number of GSVs (Fig. 6). TUG cleavage converts the upstream analog signal to a downstream digital output. A particular insulin stimulus may release a defined fraction of the pool of GSVs; if the size of the pool is larger or smaller, then the number of GSVs mobilized will be increased or decreased, respectively. Like GLUT4 (and unlike the insulin receptor, IRS and Akt), TUG proteins are not present in vast excess of those needed for a maximal response^3^. Thus, impaired Akt phosphorylation may not correlate well with impaired insulin-stimulated glucose uptake^32,40,41^. Insulin-stimulated activation of phosphatidylinositol 3-kinase (PI3K) is also reduced in insulin resistance and may act through Akt-independent targets^2,42–44^. Yet, in humans, sustained caloric restriction enhanced insulin sensitivity and, in unbiased proteomic studies of adipose tissue, increased Usp25m and reduced TUG protein abundances, with no effects on proteins that mediate PI3K signaling^32^. In rodent adipose tissue, we observed impaired TUG cleavage and reduced Usp25m abundance soon after initiation of a HFD, and prior to impaired insulin signaling through Akt^13^. More important, present data show that HFD-induced alterations occur during fasting, including dissociation of IRAP and GLUT4 targeting, increased abundance of IRAP in T-tubule membranes, and post-transcriptional depletion of Usp25m. In fasting mice, HFD feeding ablated the effect of muscle TUG knockout to cause increased glucose uptake in muscle. The data support the idea that altered Usp25m-TUG function contributes to insulin resistance, and that this occurs during fasting and independently of upstream insulin signaling.

In contrast to insulin receptor, IRS, and Akt proteins, there is minimal excess capacity of TUG proteins. In adipose and muscle, >50% of TUG proteins can be cleaved by acute, maximal insulin stimulation, at least in some experiments^13,16^. Cleavage is proportional to GLUT4 translocation, and transfers GLUT4 onto microtubule motors for trafficking to the plasma membrane^13^. Activation of microtubule-based trafficking of GLUT4 is independent of PI3K and is disrupted in insulin-resistant skeletal muscle in humans and mice^13,45,46^. In addition, the TC10 signaling pathway that triggers TUG cleavage in adipocytes may contain an upstream feedforward circuit, which could cause it to respond preferentially to the rate-of-change of plasma insulin concentration^14,28^. In muscle, the role of TC10 remains uncertain, and it is worth noting that the TC10 effector PIST may transmit signals from other Rho family members^47,48^. Nonetheless, if upstream pathways are similar to those described in adipocytes, then TUG cleavage and GLUT4 translocation might occur in proportion to glycemic load. In insulin resistant individuals, attenuation of this effect would result in postprandial hyperglycemia.

Our data support the view that distinct insulin signaling pathways, through Akt-AS160 and Usp25m-TUG, converge on the GSVs^14^. During ongoing insulin exposure, Akt-AS160 signaling may also cause GLUT4 in endosomes to return directly to the cell surface^20^. Previous data show that AS160 resides on TUG-bound, insulin-responsive GLUT4-containing vesicles, and AS160 overexpression enhances the effect of Usp25m to cause TUG cleavage in transfected cells^13^. Data here show that AS160 and TUG interact and that AS160 is required for efficient TUG cleavage in muscles stimulated *ex vivo* with insulin. These results are consistent with previous work showing that AS160 is required for the bulk of insulin-responsive GLUT4 translocation and glucose uptake in rodents^17,18,22,34,49,50^. In humans, homozygous or heterozygous null mutations in the gene encoding AS160 cause muscle insulin resistance^51,52^. Our data predict that insulin-stimulated TUG cleavage is impaired in these individuals. Much work on AS160 has focused on its activity to modulate specific Rab GTPases and, thus, to direct vesicle trafficking^24,53^. Our data imply that AS160 has functions in addition to its action on Rab proteins. Phosphorylation of AS160 disrupts its binding to IRAP and may play a role in releasing the GSVs^54^. In human muscle, the AS160 interactome includes p97 (also called VCP), an ATPase that is a major binding partner of TUG, and that is required for action of the TUG C-terminal cleavage product^16,29,55^. Understanding how these proteins act together will require further study.

To which compartments are GLUT4 and IRAP mislocalized in insulin resistant states? Data imply that GSVs bud, at least in part, from ERGIC membranes^4,12,56^. TUG localizes to the ERGIC and is ideally positioned to capture these vesicles as they are formed^29,30^. TUG may constrain a cycle of vesicle budding and fusion involving the ERGIC and possibly the *cis* cisterna of the Golgi complex. We propose that such a cycle is altered in insulin resistance, so that the pool of GSVs is depleted. In effect, the target of the insulin signal is missing. GLUT4 might be relocalized to the ERGIC or, possibly, the Golgi complex. This model is consistent with biochemical data in humans^7^. Colocalization of GLUT4 and a clathrin coat protein, CHC22, is increased in skeletal muscle from humans with type 2 diabetes, compared to nondiabetic controls^4^. CHC22 was recently shown to mediate vesicle budding at the ERGIC, and data support the idea that it is required for formation of GSVs^12,56^. CHC22 is not expressed in mice, and its function is likely performed by another clathrin, CHC17. Even so, in both mice and humans, the simplest interpretation of the data is that insulin resistance results, in part, from altered membrane trafficking causing the accumulation of GLUT4 in ERGIC membranes, as diagrammed in Fig. 6.

Our data show that HFD-induced insulin resistance is accompanied by altered targeting of IRAP in muscle, resulting in an increase in its abundance in T-tubules. A similar effect is observed in humans^7,8^ and may inform the association of hypertension with insulin resistance^57^. The formation of GSVs involves direct biochemical interactions among IRAP, GLUT4, and other proteins, resulting in ∼12 IRAP and ∼6 GLUT4 molecules in each vesicle^10,58^. These proteins are co-sequestered in the GSV pool. Loss of the co-targeting of IRAP and GLUT4 is thus a proxy for the depletion of the pool of GSVs, making the ratio of IRAP to GLUT4 in T-tubule membranes a useful surrogate. In HFD-fed mice, we observed increased targeting of IRAP to T-tubules during fasting; an even greater abundance of IRAP was observed in T-tubules after insulin stimulation. Physiologically, when present at the cell surface, IRAP cleaves and inactivates circulating vasopressin^59^. Vasopressin has a half-life of ∼1 min. in mice and, likely, in humans; this half-life is increased ∼3-fold in IRAP knockout mice and decreased ∼3-fold in mice with disruption of TUG in muscle^26,59^. Data imply that vasopressin half-life is reduced to a less marked degree in humans with insulin resistance, and that vasopressin secretion is increased as a compensatory response^57^. Copeptin, the C-terminal product of the vasopressin prohormone, is increased in the circulation of individuals with insulin resistance, and is associated with risks of diabetes, atherosclerosis, and the metabolic syndrome^60–62^. We and others have noted how chronically increased vasopressin concentrations present at the kidneys can lead to hypertension^57,63,64^. Thus, in the setting of insulin resistance, mistargeting of IRAP to T-tubule membranes may contribute directly to hypertension in humans.

We observed reduced energy expenditure in MUKO mice. This effect was more robust when data were normalized to lean mass, compared to total body mass. Upon its production, the TUG C-terminal cleavage product acts transcriptionally to enhance energy expenditure^16^. This action may contribute to the thermic effect of food^14,16^. Thus, we predicted that MUKO mice might have reduced energy expenditure resulting from impaired production of the TUG product. Like effects on glucose homeostasis, the effect of muscle Usp25m deletion to reduce energy expenditure was lost when mice were fed a HFD, consistent with reduced TUG cleavage in control mice. Previous data from MTKO mice, lacking TUG in muscle, observed reduced energy expenditure in HFD-fed mice, but not in RC-fed mice^16^. In the present work, a larger number of mice were studied on a RC diet, providing greater statistical power. It may also be that in the previous study, residual TUG cleavage in HFD-fed WT control mice was sufficient to cause a difference in energy expenditure, when compared to mice with TUG deletion. Regardless, the data support the idea that impaired TUG cleavage, as occurs in the setting of diet-induced insulin resistance, may contribute to reduced energy expenditure.

We note that Usp25m may have TUG-independent actions. A recent study described effects of Usp25 to promote the degradation of TAK1 in the heart, and thus to reduce the inflammatory response resulting from activation of JNK/p38 MAPK signaling^65^. HFD-fed mice had reduced Usp25 abundance in the heart, similar to adipose and muscle^13,16^. No effect on glucose homeostasis was observed, yet a whole body knockout was studied, hyperinsulinemic-euglycemic clamps were not performed, and no tracers were used. Knockout mice had increased HFD-induced weight gain. Thus, the phenotypes we observed overlap with those in this report; our study is arguably more sensitive because of the use of a tissue-specific knockout, and tracers to study glucose flux. Our data also raise the possibility that Usp25m acts not only to release the GSVs, but in establishment of the GSV pool itself. We observed an increased ratio of IRAP to GLUT4 in T-tubule membranes of MUKO mice. This was similar to the effect of HFD feeding and distinct from effects in MTKO mice. Therefore, it may be that Usp25m participates in the biogenesis or maintenance of the GSV pool, independent of its action to mediate TUG cleavage.

Proteases other than Usp25m may also act on TUG. In muscle lysates, we observe bands other than the 42 kDa product generated by Usp25m-mediated cleavage. It is not possible to discern whether these are produced physiologically or if they are nonspecific degradation products. Deletion of Usp25m largely eliminates the 42 kDa band. In addition to Usp25m, calpain-10 (Capn10) is implicated as a TUG protease based on studies *in vitro* and in 3T3-L1 adipocytes^66^. The relevance of Capn10 *in vivo* is not known. Capn10 may cleave TUG at several sites, distinct from the site of Usp25m cleavage, and it could function to release GLUT4 and/or to control the stability of the TUG C-terminal product. Further studies will be required to understand its physiological relevance.

Although our study focused on skeletal muscle, it is likely that a similar mechanism pertains in adipose, based on data from rodents and humans^8,13,32^. GLUT4 abundance is reduced in insulin resistant adipose, which may reflect effects on GLUT4 protein stability, as well as transcription. In 3T3-L1 adipocytes, TUG depletion enhances in GLUT4 degradation in lysosomes^19^. Thus, it is possible that reduced TUG-mediated sequestration of GSVs may contribute to reduced GLUT4 abundance in adipose in the setting of insulin resistance. Because a greater fraction of total GLUT4 normally resides in GSVs in adipose, compared to muscle, loss of GLUT4 sequestration in GSVs may have a more marked effect to decrease total GLUT4 abundance.

Mistargeting of GLUT4 and IRAP could result from altered membrane lipids, which could affect vesicle budding or fusion processes involving GSVs^2,67^. Lipids may also alter the palmitoylation of GLUT4 and other proteins, and palmitate treatment can alter GLUT4 sorting to membranes with characteristics of TUG-regulated vesicles^28,68,69^. In addition, alteration of the viscoelastic properties of a TUG-containing biomolecular condensate may control the accumulation of GSVs. In neurons, a synapsin-containing biomolecular condensate modulates the clustering of synaptic vesicles; it is possible that a similar mechanism acts in muscle to sequester GSVs^70^. In fibroblasts, deletion of TUG causes a loss of ERGIC membranes, which are absorbed into the cis-Golgi^30^. Possibly, a similar effect in muscle leads to loss of the GSV pool and impaired insulin action. Future work to define how the pool of GSVs is established and maintained will be needed to understand the pathophysiologic mechanism in detail.

In conclusion, data here support a model in which altered membrane trafficking causes the depletion of a pool of insulin-responsive vesicles, GSVs, to result in GLUT4 mistargeting and insulin resistance in skeletal muscle. This mechanism is complementary to that causing impairment of proximal steps in insulin signaling, and it likely occurs in parallel in the setting of nutritional excess. Effects to cause IRAP targeting to T-tubule membranes may contribute to hypertension, and impaired production of the TUG C-terminal cleavage product may contribute to reduced energy expenditure and worsened obesity. Thus, understanding this pathophysiology may have broad implications for the metabolic syndrome in humans.

## Methods

### Animals

Muscle Usp25m knockout (MUKO) mice were created using a targeting construct from the NIH Knockout Mouse program and the European Mouse Mutant Archive, EMMA ID: 08008, resulting in a strain carrying the floxed allele, designated C57BL/6N-A^tm1Brd^ Usp25^tm1a^(KOMP)^Mbp^/Ics. This allele contains loxP sites flanking exons 4 and 5 of the *Usp25* gene. These mice were mated with C57BL/6J mice carrying an actin-Flp transgene to remove the selection cassette, and then were backcrossed onto C57BL/6J (JAX Stock No. 000664) for at least 10 generations. To delete Usp25 in muscle, mice homozygous for the floxed Usp25 allele (Usp25^fl/fl^) were mated with Usp25^fl/+^ or Usp25^fl/fl^ mice containing a MCK-Cre transgene (Tg(Ckmm-cre)5Khn/J, JAX Stock No. 006475). Most experiments compared Usp25^fl/fl^ (WT) mice and Usp25^fl/fl^; MCK-Cre (MUKO) mice, and littermates were used as controls whenever possible. Muscle TUG knockout (MTKO) mice were described previously^16^. Male mice were used for experiments. Mice were maintained on a 12-h light–dark cycle (7 AM – 7 PM) and had ad libitum access to food and water. Mice were housed at 22 °C with 30-70% relative humidity. Most experiments used mice at age 10-15 weeks, as described. The standard regular chow (RC) diet was Harlan-Teklad 2018S and the high-fat diet (HFD) was Research Diets D12492 (60% kcal from fat). The Yale Institutional Animal Care and Use Committee approved all procedures involving mice.

For genotyping, genomic DNA was retrieved by overnight proteinase K digestion of tail or ear biopsies, and was used in PCR to assess the presence of both the floxed Usp25 allele and the MCK-Cre transgene. For the floxed Usp25 allele, PCR used 39 cycles with 95° C for 30 seconds, 59° C for 30 seconds, and 72° C for 2.5 minutes, together with the following primer pair: 5’-AGTGCCTTACAGCTCCCTCTCCTA-3’ and 5’-AAGCCTTGGTATTTGTTGATTA-3’. For the MCK-Cre transgene, PCR using a touchdown protocol was performed (JAX Protocol 23304) together with the following primer pair: 5’-GTGAAACAGCATTGCTGTCACTT-3’ and 5’-TAAGTCTGAACCCGGTCTGC-3’, and the internal positive control primer pair: 5’-CAAATGTTGCTTGTCTGGTG-3’ and 5’-GTCAGTCGAGTGCACAGTTT-3’.

AS160 knockout (KO) rats on a Wistar background were developed as previously described^34^. AS160 KO and WT control rats were maintained on a 12h light-dark cycle with ad libitum access to chow (Laboratory Diet No. 5L0D; LabDiet, St. Louis MO) and water. Male rats at 8-9 weeks of age were used for experiments. Rats were fasted beginning at 5 PM the day before the experiment, then anesthetized with an intraperitoneal injection of ketamine/xylazine cocktail (50 mg/kg ketamine and 5 mg/kg xylazine). Both epitrochlearis muscles were dissected as described^71^. Muscles were incubated for 50 min. at 35 °C in Krebs-Henseleit buffer (KHB) supplemented with 0.1% bovine serum albumin (BSA), and 2 mM sodium pyruvate, with or without 500 µU/ml insulin. Muscles were then freeze-clamped and stored at -80 °C for biochemical analyses. The University of Michigan Institutional Animal Care and Use Committee approved all procedures involving rats.

### Reagents, cell culture, transfection, and immunoprecipitation

Antibodies directed to TUG, GLUT4, and IRAP were described previously^13,19,27,37^. Other antibodies were purchased, including those directed to Usp25 (Novus NBP1-80631), GAPDH (Millipore MAB374), IRAP (Cell Signaling Technology (CST) clone D7C5, 6918S), phospho-Akt (Ser473) (CST clone D9E, 4060S), Akt (CST clone C67E7, 4691S), DHPR α-subunit (Developmental Studies Hybridoma Bank (DSHB) at Univ. of Iowa, clone IIID5E1), β-tubulin (DSHB, clone E7-b), AS160 (CST clone C69A7, 2670S), and anti-Flag M2 (MilliporeSigma F3165). Anti-Flag M2 affinity gel (MilliporeSigma A2220) was used for immunoprecipitations. For immunoblots, primary antibodies were used at 1:1000 dilution or at ∼2 μg/ml unless otherwise noted. Total protein was assessed using a Pierce Reversible Protein Stain Kit for Nitrocellulose Membranes (ThermoFisher Cat. No. 24580).

HEK293 cells were cultured in high glucose DMEM GlutaMAX medium (Invitrogen) containing 10% EquaFETAL bioequivalent serum (Atlas Biologicals), antibiotic-antimycotic solution (Sigma), and plasmocin (Invivogen). Lipofectamine 2000 (Invitrogen) was used for transient transfection according to the manufacturer’s protocol. Plasmids to express flag-tagged AS160 and flag-tagged TNKS2 constructs were described previously^72–74^ and were gifts of Dr. Gustav Lienhard and Dr. Nai-Wen Chi, respectively. The flag-TNKS2-ANK plasmid contains only the ankyrin repeat domain of TNKS2 (residues 1-182 of the intact 1166 residue protein). TUG plasmids were described previously^37^.

For immunoprecipitation, cells were lysed 48 h after transfection in buffer containing 0.5% IGEPAL CA-630 (Sigma), 20 mM Tris, pH 7.4, 150 mM NaCl, 1 mM EDTA with Complete (Roche) protease inhibitors. After centrifugation to clear insoluble debris, an aliquot was saved for control immunoblots and lysates were incubated overnight with anti-Flag M2 affinity gel (above) at 4 °C with end-over-end rotation. The next day, beads were pelleted in a benchtop microfuge, washed 6 times with 0.5% IGEPAL CA-630 buffer and transferred to new tubes. Samples were eluted by heating (5 min., 95° C) in SDS-PAGE sample buffer with 15% 2-mercaptoethanol. Eluates were separated on 4-12% NuPAGE bis-tris gels (Invitrogen) and immunoblotted to nitrocellulose membranes using a semi-dry transfer apparatus (Biorad).

### Biochemical analyses and immunoblotting

To isolate basal or insulin-stimulated tissues from mice, WT or MUKO mice were fasted for 6 hours, then treated with intraperitoneal (IP) injection of insulin (10 U/kg) and glucose (2 g/kg) in phosphate buffered saline (PBS), or with an equivalent volume (∼0.3 ml) of PBS alone, as previously^16^. After 30 min., mice were anesthetized and sacrificed by cervical dislocation. Tissues were collected and flash frozen in liquid nitrogen and stored at -80° C.

To isolate T-tubule-enriched membrane fractions, we used quadriceps muscles, as previously described^16,38^. Briefly, ∼12-week old mice were fasted 6 h, then treated by IP injection of insulin and glucose solution, or saline control, as above. After 30 min., mice were anesthetized and euthanized by cervical dislocation. One entire quadriceps muscle (∼100 mg) was trimmed, minced in 2 ml of ice-cold Buffer A (20 mM Na_4_P_2_O_7_, 20 mM NaH_2_PO_4_, 1 mM MgCl_2_, 0.3 M sucrose, 0.5 mM EDTA, 20 mM iodoacetamide, and two protease inhibitor tablets (Roche) per 50 ml). Samples were homogenized using a Polytron tissue grinder for 10-15 sec. at 8000 rpm and then centrifuged at 13,000 rpm for 20 min. at 4 °C using a SS-34 rotor (Sorvall). Pellets were resuspended in 1.5 ml of Buffer A, homogenized again using a Polytron (13,500 rpm for 45 sec.), and then centrifuged at 11,000 rpm for 20 min. at 4 °C using an SS-34 rotor. The supernatant was placed in a new tube and centrifuged in a TLA-120.2 rotor (Beckman) at 18,000 rpm for 10 min. at 4 °C. Supernatant was removed carefully and pellets were resuspended in 200 µL of Buffer A. To strip myofibrillar proteins, 500 µl of Buffer B (0.3 M sucrose, 20 mM Tris, pH 7.0) and 300 µl of 4M KCl were added to make a final KCl concentration of 1.2 M. Samples were incubated at 4 °C for 1 h with gentle agitation. Samples were then centrifuged at 57,000 rpm in a TLA-120.2 rotor for 10 min at 4 °C. The pellets representing the T-tubule-enriched membrane fraction were resuspended in 150 µl of Buffer B, mixed with 4x LDS loading buffer (Invitrogen) and analyzed by SDS-PAGE and immunoblotting.

For immunoblots of tissue lysates, tissues were quickly thawed and 30-50 mg of each tissue were weighed and mixed with lysis buffer (1% IGEPAL CA-630 (Sigma), 20 mM Tris, pH 7.4, 150 mM NaCl, 1 mM EDTA with Complete (Roche) protease inhibitors). A Qiagen TissueLyser II was used to grind the tissue for 1.5 min. at 30 cycles/sec. Samples were kept on ice for 20 min. to allow lysis to proceed, with vortexing every 5 min. To remove insoluble debris, lysates were centrifuged 10 min at 13,000 rpm in a tabletop centrifuge (Eppendorf 5424R) at 4° C. Supernatants were analyzed by SDS-PAGE using Invitrogen NuPAGE gels, transferred to nitrocellulose membranes using a semidry apparatus, and imaged using peroxidase conjugated secondary antibodies and detection on film, as previously^13,16^. Quantification was done on exposures within the linear range of the film and used transillumination (Epson Perfection V700 flatbed scanner) together with Silverfast 9 (Lasersoft Imaging, version 8.8.0) and ImageJ (FIJI, version 2.1.0/1.53c) software. Figures were prepared using Adobe Photoshop (version 25.12.0) and Illustrator (version 28.7.1; Creative Cloud 2024), and Graphpad Prism (version 10.0 for MacOS) software.

### Metabolic analyses

Fasting blood glucose and insulin concentrations were measured using a handheld glucometer (Zoetis AlphaTrak3) and by an ultrasensitive ELISA (Mercodia 10-1249-01). For tail vein glucose measurements, mice were fasted for 4 h in separate cages with unrestricted access to water. Mice were restrained by hand, and an 18 g needle was used to create a single venipuncture site on the tail vein. Blood was expressed through manual milking of the tail and one large drop (∼10 µl) was applied to a glucometer. HOMA-IR was calculated as [glucose]*[insulin] / 22.5, where glucose is given in mmol/l and insulin is in mU/l. Plasma NEFA was measured using the enzymatic colorimetric method by Wako reagents (FUJIFILM; Wako Diagnostics, USA).

Fat and lean mass were measured using ^1^H NMR (Minispec, Bruker Biospin^15,16^. Rates of oxygen consumption (VO_2_) and carbon dioxide production (VCO_2_), energy expenditure, respiratory exchange ratio, locomotor activity, food consumption, and water intake were measured using CLAMS metabolic cages (Columbus Instruments), as previously^15,16^, and were analyzed as described^75^.

For diet studies, WT and MUKO mice were fed a HFD beginning at 8 weeks of age, and body weights were measured twice weekly. Mice were studied for 7 weeks on the HFD, then euthanized. GWAT was isolated and weighed.

Dynamic measurements of glucose flux were done as previously, using a Xylem YSI 2500 Biochemistry Analyzer to measure glucose concentrations^15,16,76,77^. Briefly, a catheter was placed in the right jugular vein 6-7 days before turnover studies were done. Only mice that regained body weight to that prior to this procedure were studied. After an overnight fast, 3-[^3^H]glucose (Perkin-Elmer Life Sciences) was infused at a rate of 0.05 μCi/min for 120 min for basal glucose turnover measurement. A blood sample was collected by tail bleeding at the end of the basal infusion to determine plasma glucose, insulin, and NEFA concentrations and ^3^H-glucose specific activity. Following the basal infusion, a 140 min hyperinsulinemic-euglycemic clamp was performed with a continuous infusion of insulin at the dose of 2 mU kg^−1^ min^−1^ in RC-fed mice, or at 3 mU kg^−1^ min^−1^ in HFD-fed mice. Dextrose (20%) was infused at variable rates to maintain euglycemia (110-130 mg/dl). The dextrose was enriched with [3-^3^H]glucose to measure insulin-stimulated glucose turnover. A bolus (10 μCi) of 2-[1-^14^C]deoxyglucose (2-DOG) was injected at 85 min to determine the rate of muscle-specific glucose uptake under insulin stimulation. During the clamp, blood samples were collected at 10–25 min intervals for immediate determination of plasma glucose levels. After 2-DOG injection, plasma was collected to measure plasma ^14^C-glucose specific activity. At the end of the clamp, plasma was collected to measure plasma glucose, insulin, and NEFA concentrations and ^3^H/^14^C-glucose specific activity.

For measurement of whole body and muscle-specific glucose uptake during fasting, MTKO and WT control mice were fed a HFD beginning at 8 weeks of age. Jugular vein catheters were placed 19 days after beginning the HFD, and turnover studies were done 26 days after beginning the HFD. Fasting glucose turnover studies were done after a 6 h fast, as previously^16^. The protocol was similar to that for the hyperinsulinemic-euglycemic clamp studies described above, except that insulin was not infused and mice were monitored closely to minimize any increases in plasma glucose concentrations. To measure muscle-specific fasting glucose uptake, 10 μCi of 2-deoxy-D-[1-^14^C] glucose was infused, without insulin and with monitoring of plasma glucose and care to minimize any increases. Blood samples were drawn from the tail vein at 5, 15, 25, 35, 45, and 55 min after 2-deoxyglucose infusion. At the end of each study, mice were treated with intravenous pentobarbital sodium injection (150 mg/kg), tissues were quickly excised, snap frozen in liquid nitrogen, and stored at -80° C for subsequent analysis. Intracellular (6-phosphorylated) 2-deoxyglucose was measured and used to calculate tissue-specific glucose transport as described previously^15,16,76,77^.

### Quantitative real time PCR (qPCR) Analysis

Total RNA was prepared from quadriceps muscles of WT and MUKO mice using Trizol reagent. Briefly, tissues were homogenized in 1 ml Trizol solution (ThermoFisher) using a Qiagen TissueLyser II at 30 cycles/sec for 1.5 minutes. After homogenization, 0.2 ml chloroform was added. After vigorous vortexing, the mixture was centrifuged at 12,000 rpm in a benchtop centrifuge (Eppendorf 5424R) at 4° C for 15 min. The aqueous phase was transferred to a new tube, and RNA was precipitated by adding 0.5 ml isopropyl alcohol. RNA was pelleted by centrifugation at 12,000 rpm at 4°C for 10 min. in a benchtop centrifuge, washed with 75% ethanol, and then centrifuged again at 7,500 rpm at 4°C for 5 minutes. After drying, total RNA was redissolved in RNase-free water. After nanodrop concentration, RNA was stored in -80°C for future use. To synthesize cDNA, ∼1 µg total RNA was used, together with a High-Capacity cDNA Reverse Transcription Kit (Applied Biosystems). cDNA was stored at -20°C. Real-time qPCR was performed using iTaq Universal SYBR Green Supermix (Bio-Rad) on a StepOnePlus Real Time PCR system (Applied Biosystems) following the manufacturer’s instructions. To detect Usp25 transcript abundance, primers were: 5’-TCCGGCACCAAGGCACATCAC-3’ and 5’-ACGGCATGGAGGCGGTAAGG-3’. Data were normalized to β-actin, using primers 5’-TGGAATCCTGTGGCATCCATGAAAC-3’ and 5’-TAAAACGCAGCTCAGTAACAGTCCG-3’. Relative abundance of Usp25 transcripts was calculated by the Λ1Λ1Ct method.

### Statistics and Reproducibility

All data were replicated in at least two independent experiments, and usually three or more replicates were performed using biologically independent samples. On bar graphs showing individual replicates, data are presented as mean ±S.E.M. Biological replicates indicate that data were obtained using different mice, different muscles ex vivo, or different plates of cultured cells. Significance was assessed using unpaired, two-tailed t-tests or using one-way ANOVA with Bonferroni adjustment for multiple comparisons of preselected pairs. Exact p values are indicated in the figures, except for p<0.0001. Differences were considered significant at p<0.05. No statistical tool was used to pre-determine sample sizes; rather, the availability of materials and estimates of variances based on previous experience determined the number of biological replicates that were used.

## Acknowledgements

We thank Estifanos Habtemichael, Anup Parchure, Daniel Vatner, William Philbrick, Pamela Dann, Gustav Lienhard, and Nai-Wen Chi for advice, reagents, and assistance. This work used Core Facilities of the Yale Diabetes Research Center (DRC, NIH P30 DK045735) and the Yale Mouse Metabolic Phenotyping Center Live (U2C DK134901). This work was supported by NIH grant R01 DK129466, the Blavatnik Family Foundation, and a Yale POINTs Pilot Award (to J.S.B). G.D.C acknowledges support from NIH R01 DK136700. G.I.S. acknowledges support from NIH RC2 DK120534 and R01DK133143).

## Supplementary Figure Legends

**Suppl. Fig. 1.**
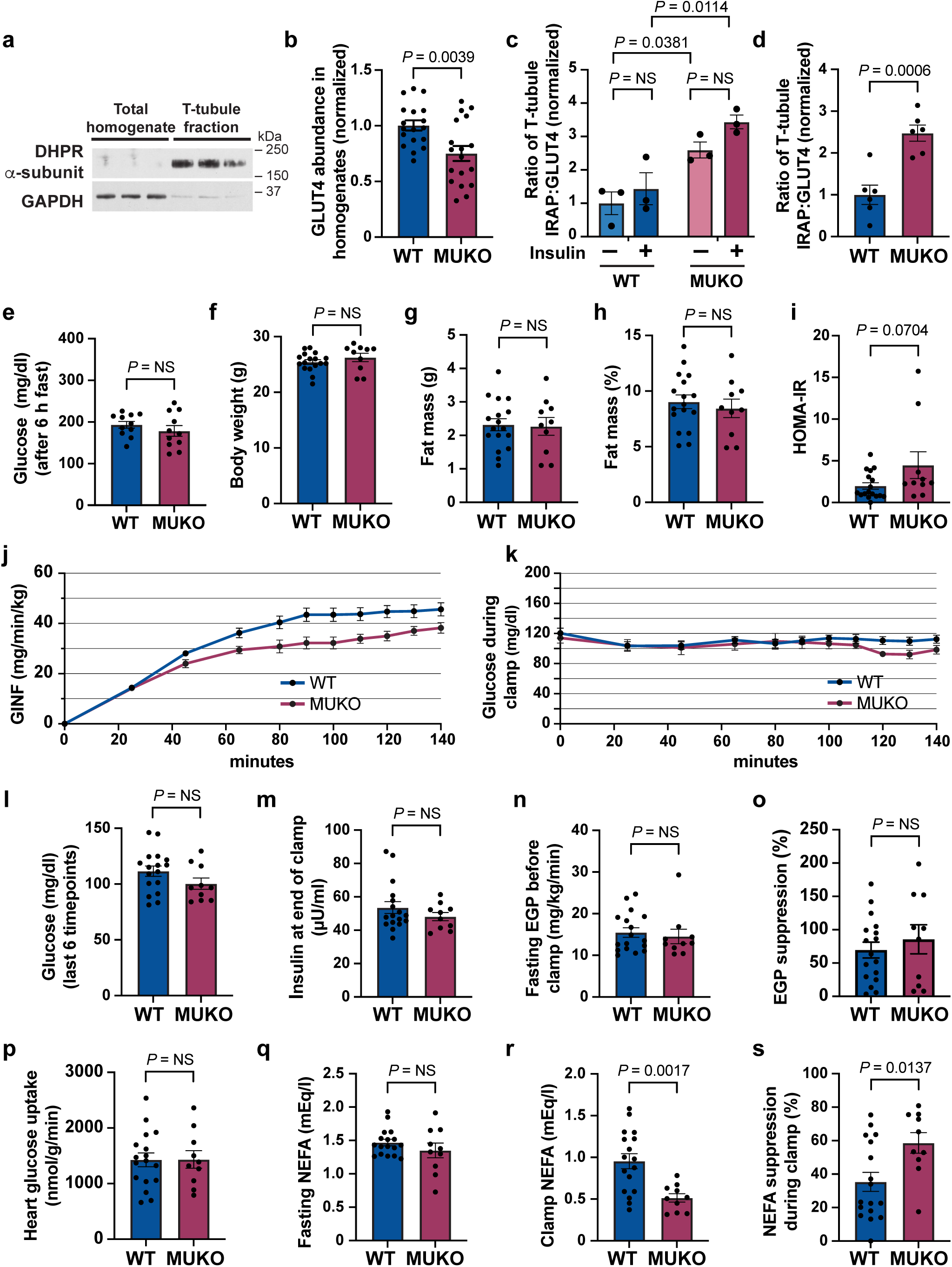
Supporting data for studies of glucose turnover in MUKO mice. **a.** Quadriceps muscles were isolated from WT mice, homogenized, and T-tubule enriched membrane fractions were prepared. Immunoblots were performed as indicated to demonstrate the purity of the fractions. **b.** GLUT4 was immunoblotted in total homogenates of WT and MUKO mice (Fig. 1c), data were quantified and are plotted. n = 18 mice in each group. **c. ,d.** IRAP and GLUT4 abundances were quantified in T-tubule fractions of WT and MUKO mice (Fig. 1c-e), and the ratio of IRAP to GLUT4 abundance is plotted. Data are plotted for fasted and insulin-stimulated mice (c) and in all WT and MUKO mice (d). In (c), n = 3 mice in each group. In (d), n = 6 mice in each group. NS, not significant. **e.** Plasma glucose concentrations were measured in 12-week-old WT and MUKO mice fasted for 6 h. n = 11 mice in each group. NS, not significant. **f.-h.** Body weight (f), fat mass (g), and percent fat mass (j) were measured in 12-week-old mice used for hyperinsulinemic-euglycemic clamps. n = 17 WT and 10 MUKO mice. NS, not significant. **i.** HOMA-IR was calculated from fasting glucose and insulin concentrations measured prior to hyperinsulinemic-euglycemic clamps. n = 17 WT and 10 MUKO mice. **j. ,k.** Glucose infusion rate (GINF) and plasma glucose concentrations are plotted versus time during the hyperinsulinemic-euglycemic clamp. Mean ±s.e.m., n = 17 WT and 10 MUKO mice. **l.-s.** The indicated parameters were measured in hyperinsulinemic-euglycemic clamps. n = 17 WT and 10 MUKO mice. EGP, endogenous glucose production. NEFA, nonesterified fatty acid. NS, not significant. All data are presented as mean ± s.e.m. of biologically independent samples, analyzed using a two-tailed t test (c-h,k-r) or ANOVA with adjustment for multiple comparisons (b).

**Suppl. Fig. 2.**
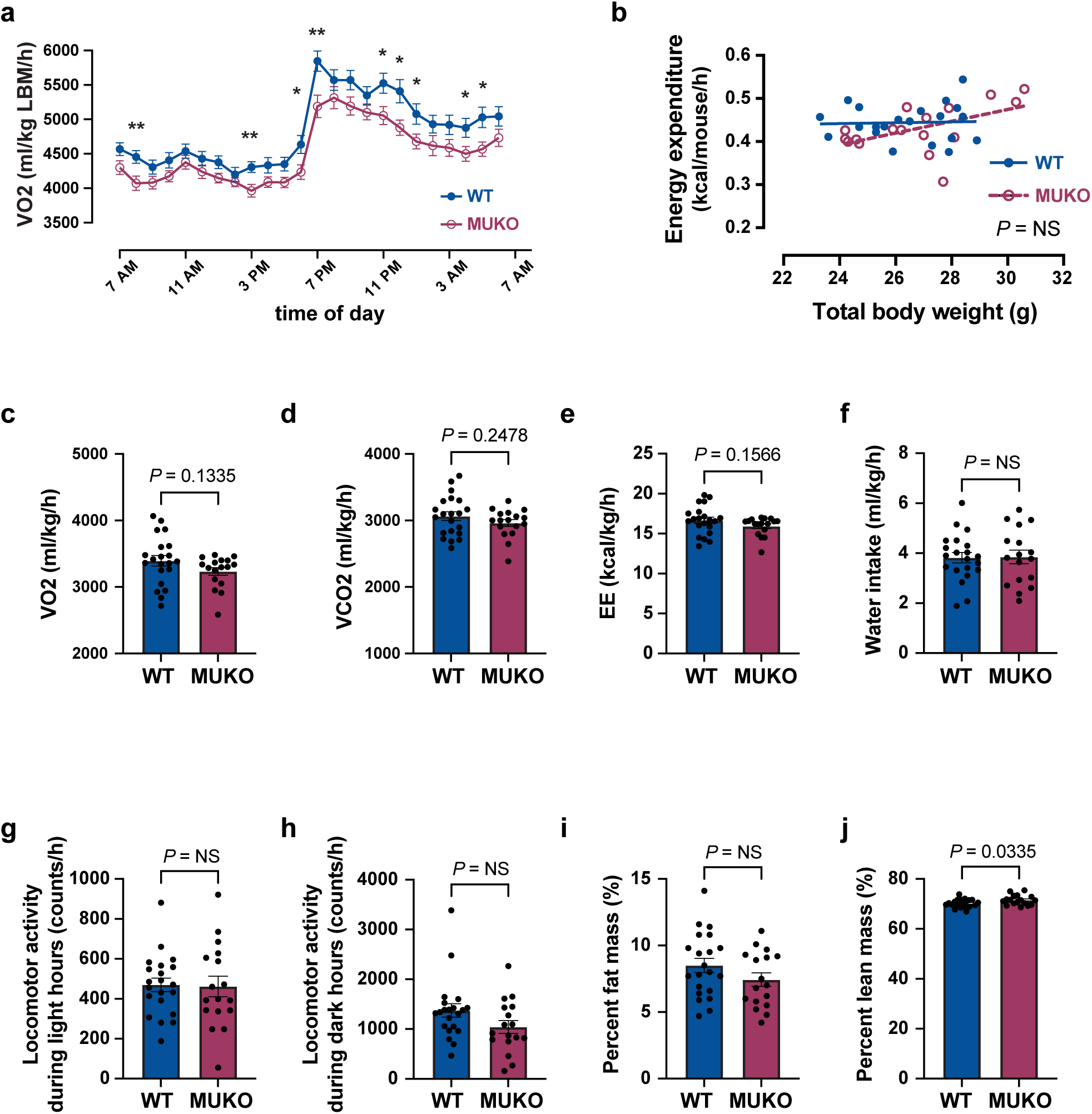
Supporting data for studies of energy expenditure in MUKO mice. **a.** Rate of oxygen consumption (VO_2_) was measured by in metabolic cages in 13-week-old mice and is plotted over time. n = 21 WT and 17 MUKO mice. Mean ± s.e.m. is shown. Individual time points were analyzed using two-tailed t tests. \**P*<0.05, \*\**P*<0.01. **b.** Energy expenditure per mouse is plotted as a linear regression versus total body weight. n = 21 WT and 17 MUKO mice. Data were analyzed by ANCOVA. NS, not significant. **c.-j.** The indicated parameters were measured in metabolic cages (c-h) or in body composition analyses done immediately before metabolic cage analyses. n = 21 WT and 17 MUKO mice. Data are presented as mean ± s.e.m. of biologically independent samples, analyzed using a two-tailed t test. NS, not significant.

**Suppl. Fig. 3.**
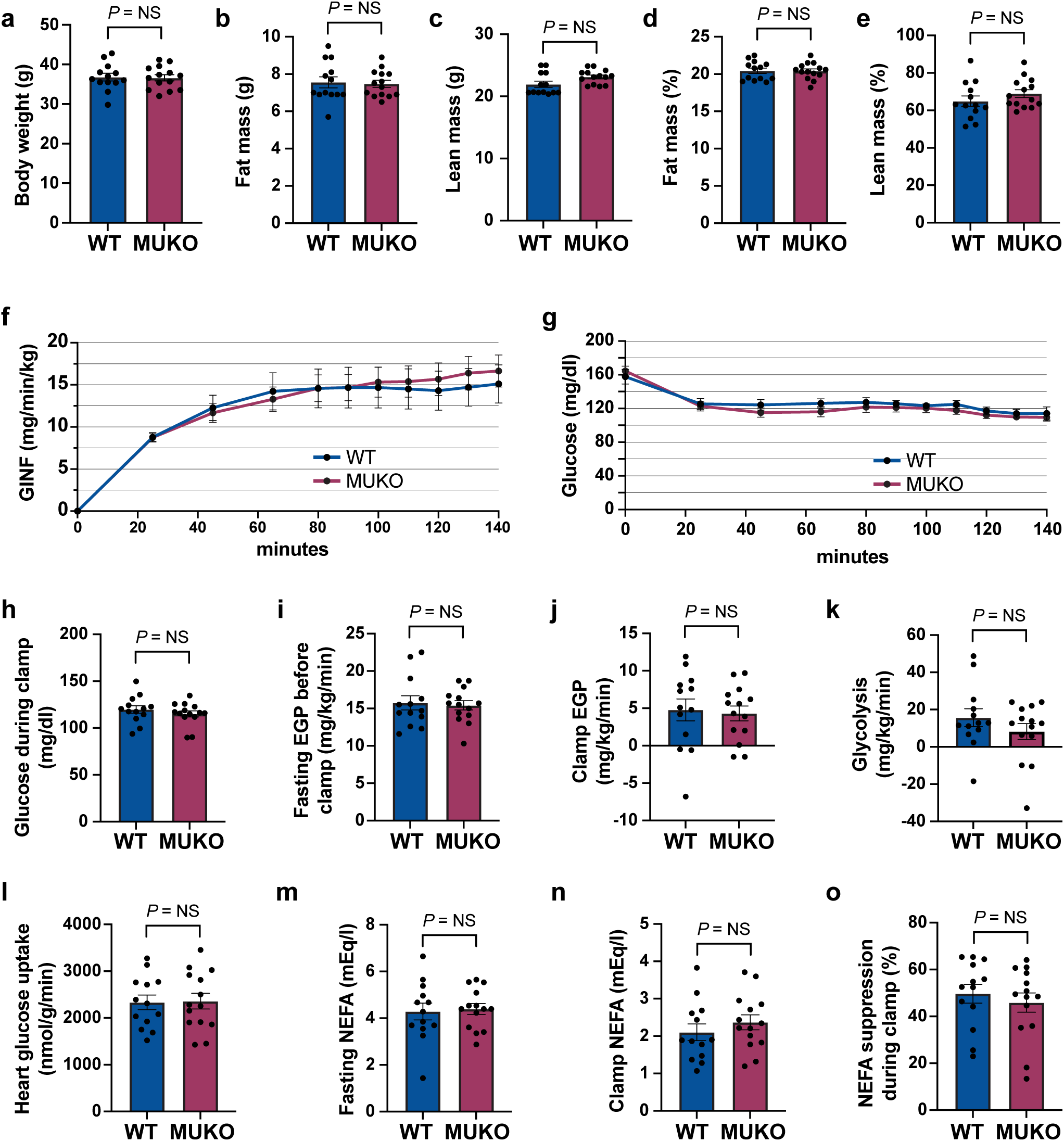
Supporting data for studies of glucose turnover in HFD-fed MUKO mice. **a.-e.** Body weight and composition were measured prior to hyperinsulinemic-euglycemic clamp studies, and the indicated parameters are plotted. n = 13 WT and 14 MUKO mice. Mean ± s.e.m. is shown, analyzed using two-tailed t tests. NS, not significant. **f.,g.** Glucose infusion rate (GINF) and plasma glucose concentrations are plotted versus time during the hyperinsulinemic-euglycemic clamp. Mean ±s.e.m., n = 13 WT and 14 MUKO mice. **h.-o.** The indicated parameters were measured in hyperinsulinemic-euglycemic clamps. n = 13 WT and 14 MUKO mice. Data are presented as mean ± s.e.m. of biologically independent samples, analyzed using a two-tailed t test. EGP, endogenous glucose production. NEFA, nonesterified fatty acid. NS, not significant.

**Suppl. Fig. 4.**
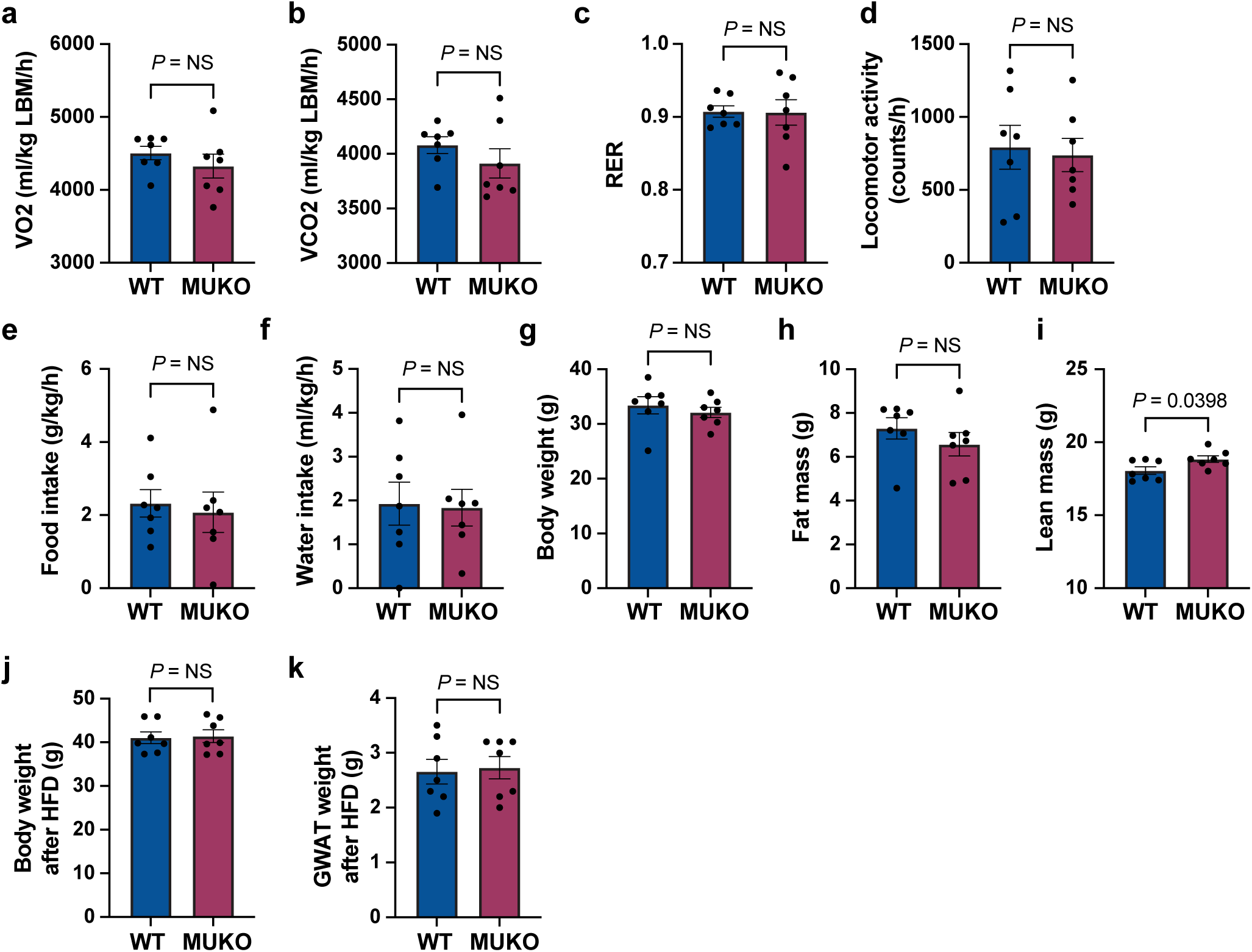
Supporting data for studies of energy expenditure in HFD-fed MUKO mice. **a.-i.** Mice were treated with a HFD for 3 weeks, then studied at age 12 weeks using indirect calorimetry and analysis of body weight and composition. The indicated parameters were measured and are plotted. n = 7 WT and 9 MUKO mice. **j,k.** Mice were fed a HFD beginning at 8 weeks of age. Body weight (j) and GWAT weight (k) were measured after 7 weeks. n = 7 mice in each group. All data are shown as mean ± s.e.m., analyzed using a two-tailed t test. NS, not significant.

**Suppl. Fig. 5.**
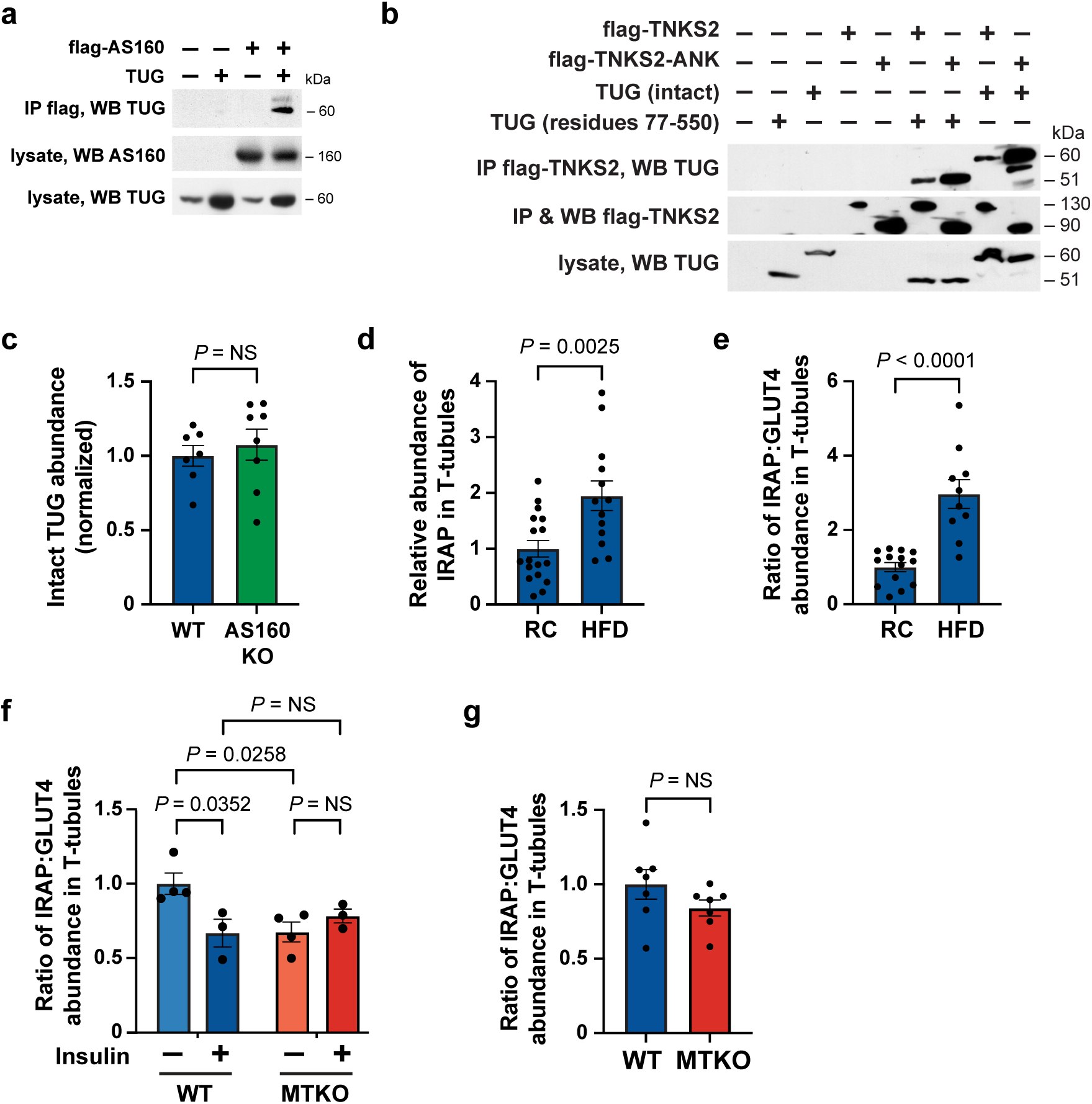
Supporting data to characterize GLUT4 Storage Vesicle -regulating proteins. **a.** TUG and flag-tagged AS160 were expressed by transient transfection of HEK293 cells. Cells were lysed and flag-AS160 was immunoprecipitated. Eluates and lysates were immunoblotted as indicated. IP, immunoprecipitate. WB, western blot. **b.** TUG and flag-tagged TNKS2 proteins were expressed by transient transfection of HEK293 cells. The flag-TNKS2-ANK construct contains only the ankyrin repeat domain of TNKS2. Cells were lysed and flag-TNKS2 proteins were immunoprecipitated. Eluates and lysates were immunoblotted as indicated. IP, immunoprecipitate. WB, western blot. **c.** Lysates from epitrochlearis muscles of WT and AS160 KO rats were immunoblotted for TUG. Data were quantified and normalized to that in WT controls. n = 7 WT and 8 KO samples. **d. ,e.** T-tubule membrane fractions were isolated from quadriceps muscles of 14-week-old mice that had been maintained on regular chow (RC), or fed a high-fat diet (HFD) for 6 weeks, as described for Fig. 4h-k. As similar numbers of saline-and insulin-treated samples were analyzed in each group, the overall abundance of IRAP (d) and ratio of IRAP to GLUT4 (e) were quantified by genotype. For (d), n = 17 RC and 13 HFD samples. For (e), n = 14 RC and 10 HFD samples. **f.,g.** T-tubule membrane fractions were previously isolated from saline-and insulin-treated WT and muscle TUG knockout (MTKO) mice, and were immunoblotted to detect IRAP and GLUT4^16^. These previous data were analyzed to quantify the ratio of IRAP to GLUT4 (f). n = 4 saline WT, 3 insulin WT, 4 saline MTKO, and 3 insulin MTKO samples. In (g), data were grouped by genotype. All data are presented as mean ± s.e.m. of biologically independent samples, analyzed using a two-tailed t test (c,d,e,g) or ANOVA (f). NS, not significant.

**Suppl. Fig. 6.**
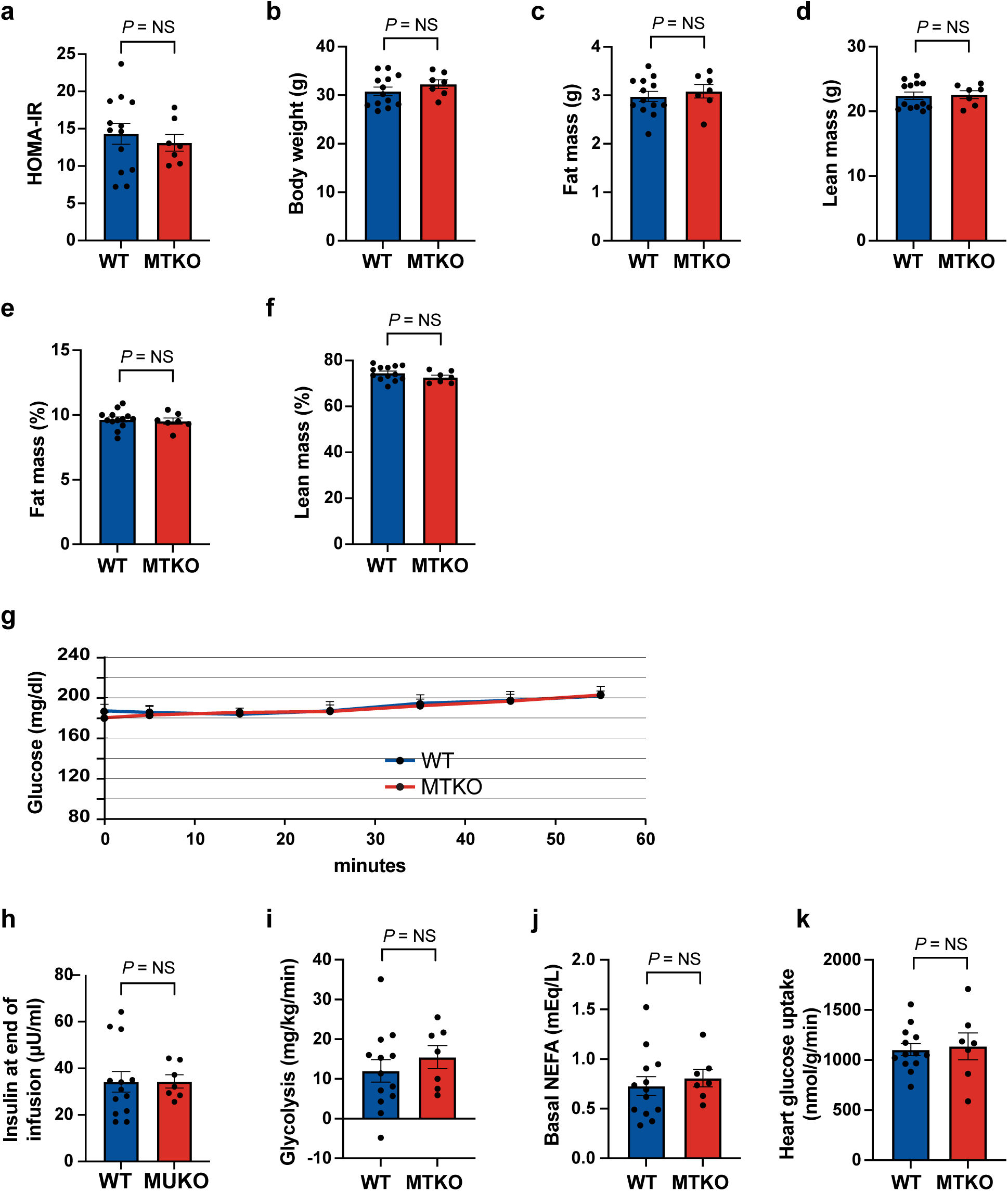
Supporting data for studies of glucose turnover during fasting in high-fat diet -fed muscle-specific TUG knockout mice. **a.** HOMA-IR was calculated from 6-h fasted glucose and insulin concentrations prior to glucose turnover studies presented in Fig. 5. n = 13 WT and 7 MTKO mice. **b.-f.** Body weight and composition were measured and the indicated parameters are plotted. n = 13 WT and 7 MTKO mice. **g.** Glucose concentrations in HFD-fed WT and MTKO mice during the fasting glucose turnover study. n = 13 WT and 7 MTKO mice. mean ±s.e.m. is plotted. **h.-k.** The indicated parameters were measured in HFD-fed WT and MTKO mice during the fasting glucose turnover study. n = 13 WT and 7 MTKO mice. NEFA, nonesterified fatty acid All data are presented as mean ± s.e.m. of biologically independent samples, analyzed using a two-tailed t test. NS, not significant.

